# External fertilization is orchestrated by a pH-regulated soluble adenylyl cyclase controlling sperm motility and chemotaxis

**DOI:** 10.1101/2021.06.18.448929

**Authors:** H.G. Körschen, H. Hamzeh, R. Pascal, L. Alvarez, W. Bönigk, N. Kaur, L.R. Levin, J. Buck, C. Kambach, M. Michino, A. Jennings, A. Sato, R. Seifert, T. Strünker, C. Steegborn, U.B. Kaupp

## Abstract

The reaction of CO_2_ with H_2_O to form HCO_3_^-^ and H^+^ is one of the most important chemical equilibria in cells. In mammalian sperm, a soluble adenylyl cyclase (sAC) serves as cellular HCO_3_^-^ sensor that conveys the equilibrium state via cAMP synthesis to cAMP-signaling molecules. The function of sAC and cAMP in non-mammalian sperm is largely unknown. Here, we identify sAC orthologs in sea urchin and salmon sperm that, surprisingly, are activated by alkaline pH rather than HCO_3_^-^. Two amino-acid residues required for HCO_3_^-^ binding of mammalian sAC are lacking in pH-regulated sAC. Orthologs identified in ten other phyla are also lacking either one of these key residues, suggesting that pH control is widespread among non-mammalian metazoan. The pH-sensitive sAC controls several functions of sperm from external fertilizers. Upon spawning, alkalization triggers cAMP synthesis and, thereby, activates motility of quiescent sperm. Egg-derived chemoattractants also alkalize sperm and elevate cAMP, which then-modulates pacemaker HCN channels to trigger a chemotactic Ca^2+^ response. Finally, the sAC and the voltage- and cAMP-activated Na^+^/H^+^ exchanger sNHE mutually control each other. A picture of evolutionary significance is emerging: motility and sensory signaling of sperm from both internal and external fertilizers rely on cAMP, yet, their sAC is regulated by HCO_3_^-^ or pH_i_, respectively. Acidification of aquatic habitats due to climate change may adversely affect pH-sensing by sAC and thereby sexual reproduction in the sea.

**Statement of significance:** Adenylyl cyclases synthesize cAMP, a prominent cellular messenger. A bicarbonate-sensitive AC family member, soluble AC (sAC), is tied to the chemical equilibrium: H_2_O + CO_2_ ↔ HCO_3_^-^ (bicarbonate) + H^+^. The sAC is required for fertilization: Mammals lacking sAC are infertile and sperm immotile. We now identify a new sAC form in sperm of non-mammalian animals that reproduce in the sea. This novel sAC is activated at alkaline pH rather than bicarbonate. It controls sperm motility and chemotaxis. The switch from HCO_3_^-^ to pH rests on substitution of two amino-acids, which represents an adaptation to aquatic environments low in bicarbonate. Acidification of aquatic habitats due to climate change may adversely affect sAC activity and, thereby, fertilization.

## Introduction

Cyclic AMP is a ubiquitous cellular messenger that orchestrates many physiological processes. In metazoans, it is synthesized by two different types of adenylyl cyclases: transmembrane AC (tmAC) and soluble AC (sAC). The sACs orthologs studied so far are directly regulated by bicarbonate (HCO_3_^-^) and calcium (Ca^2+^) (1). Cellular HCO_3_^-^ levels are determined by the chemical equilibrium:

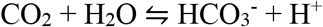

The sAC monitors the status of this equilibrium and translates changes into a cAMP signal that controls downstream targets directly or via phosphorylation by protein kinase A (PKA) (1). Its key role as HCO_3_^-^ sensor has been firmly established in mammalian sperm. Soluble AC may also serve as pH_i_ sensor indirectly via the CO_2_/HCO_3_^-^/H^+^ equilibrium (1). The HCO_3_^-^ concentration (15-25 mM) in semen and the oviduct stimulates cAMP synthesis by sAC, which initiates and modulates sperm motility. Moreover, cAMP synthesis by sAC initiates an essential maturation process, called capacitation (2). Deletion of the genes encoding sAC or the regulatory and catalytic subunits of PKA renders sperm immotile and male mice infertile (3-5). In humans, frame-shift mutations in the *adcy10* gene encoding sAC also cause male infertility due to immotile sperm (6).

Soluble AC and potential cAMP-signaling targets have also been identified in sperm of external fertilizers ranging from marine invertebrates to fish (7-12). Although its properties seem to be conserved across phyla (7, 13, 14), the role of sAC in sperm exposed to aquatic habitats low in HCO_3_^-^ (2-4 mM) is enigmatic.

Here, we show that sAC from sea urchin and fish is directly activated at alkaline pH but not by HCO_3_^-^ and Ca^2+^, connecting cAMP synthesis directly with pH_i_ homeostasis. We define the role of sAC and cAMP for motility activation and chemotactic signaling. Substitutions of two amino-acid residues in the HCO_3_^-^-binding site of mammalian sAC, which predicts the switch from HCO_3_^-^ to pH regulation, are conserved across ten phyla, suggesting that the pH regulation is conserved across non-mammalian sperm.

## Results

### Mammalian and non-mammalian sAC differ in two key amino acids

The crystal structure of the human sAC catalytic core (sAC-cat) revealed key amino acids involved in binding of HCO_3_^-^ and ATP/ion substrates (15-18). In non-mammalian sAC, most of these residues are conserved (Supplementary Table 1) except for positively charged R176 and K95 that coordinate the HCO_3_^-^ anion. In 15 out of 17 marine invertebrates examined, R176 is replaced by Asn, Ser, or Thr, while K95 is conserved (Supplementary Table 2). In both bony and cartilaginous fish (again, with one exception, Coelacanthiformes *L. chalumnae)*, K95 is replaced by Asn or Thr, while R176 is conserved (Supplementary Table 2). Thus, the sAC of most aquatic animals lacks either one of these two key amino-acid residues that endow mammalian sAC with HCO_3_^-^ sensitivity.

We studied the structural consequences in a homology model of the catalytic core of the *s*AC from *A. punctulata* (*Ap*sAC-cat), consisting of two catalytic domains C1 (residues 65-229) and C2 (residues 338-542). Sequence comparison revealed that the cyanobacterial sAC homologue CyaC (PDB ID 1WCO (19)) and C2 from human sAC (PDP ID 4CLL (15)) display the highest sequence similarity to *Ap*sAC-cat C1 and C2, respectively. Use of these two templates yielded an excellent homology model for *Ap*sAC-cat (Fig. 1A) with proper positioning of the conserved catalytic residues (15). K117 is positioned toward HCO_3_^-^ in the regulatory site of *Ap*sAC, similar to K95 in human sAC (Fig. 1B). The smaller N198, which corresponds to human R176, is also oriented toward HCO_3_^-^; however, N198 remains more distant due to its shorter sidechain and a conserved main-chain fold in this region. Interestingly, the shorter sidechain results in an opening that renders the regulatory site accessible via a channel from the protein surface (Fig. 1C). In human sAC, R176 blocks this channel and access to the regulatory site is solely provided by a channel that branches off the active site. This analysis suggests that non-mammalian sAC activity may not rely on HCO_3_^-^.

**Figure 1.**
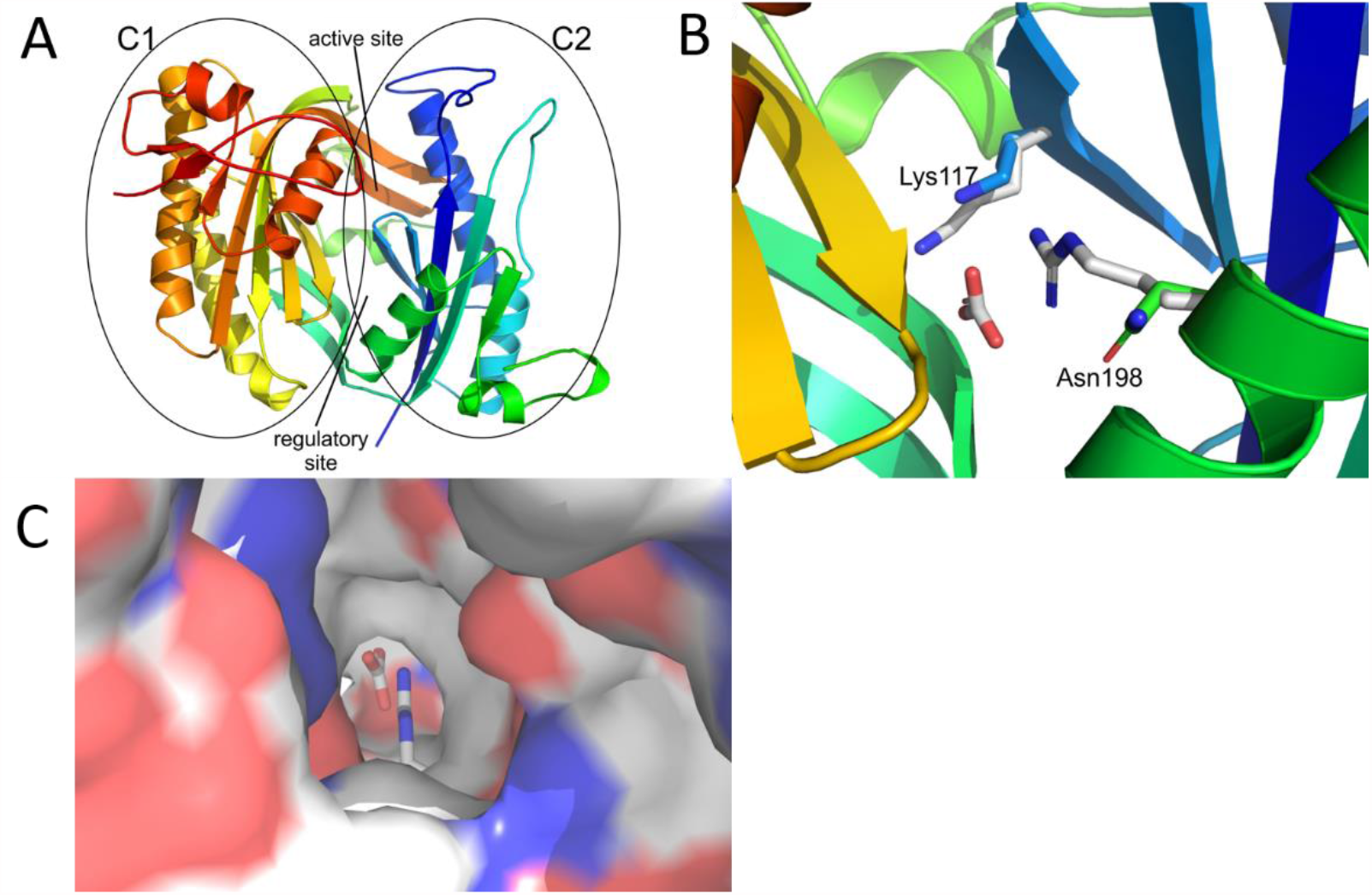
Homology modeling of the soluble adenylyl cyclase sAC from *A. punctulata*. **(A)** Overall structure of the catalytic cores C1 and C2, in rainbow coloring from N-terminus (blue) to C-terminus (red). C1, C2 as well as the active site and the pseudo-symmetric regulatory site are indicated. **(B)** Regulatory site of *Ap*sAC-cat lined by K117 (blue) and N198 (green) and overlayed with the corresponding human sAC residues binding bicarbonate (grey sticks). **(C)** View from the protein surface into the regulatory site of *Ap*sAC colored according to electrostatic potential (negative red, positive blue) and overlayed with human sAC (sticks). The view shows an access channel in *Ap*sAC that is blocked by R176 in human sAC.

### The sAC of sea urchin and salmon is regulated by pH but not HCO_3_^-^ and Ca^2+^

We chose sAC orthologues from *A. punctulata* (*Ap*sAC) and salmon *S. salar* (*Ss*sAC) to study the functional consequences of R176N and K95N substitutions. Mammalian sAC, e.g., from rat (*Rn*sAC), exists as full-length (fl) and truncated (t) isoforms (sAC_fl_, 180 kDa and sAC_t_, 55 kDa, respectively) (20, 21). The specific activity of sAC_t_ is much greater than that of sAC_fl_ due to an autoinhibitory domain C-terminal to the C2 catalytic domain (22) that is conserved in mammalian and non-mammalian sACs. *A. punctulata* sperm contain only sAC_fl_ (7, 9). To allow for direct comparison with rat *Rn*sAC_t_, we studied artificially truncated *Ap*sAC and *Ss*sAC. *Ap*sAC, *Ss*sAC, and *Rn*sAC were heterologously expressed in HEK293 cells, and cAMP synthesis was determined by ELISA.

Because [HCO_3_^-^] in water is low compared to tissue levels, we stimulated sAC at various [HCO_3_^-^]. Unlike mammalian sAC, the activity of *Ap*sAC_t_ and *Ap*sAC_fl_ was not or only weakly affected by [HCO_3_^-^] at both neutral and alkaline pH (Fig. 2A, B). Moreover, cAMP synthesis of *Ap*sAC_t_ (MgATP as substrate) was not significantly affected by [Ca^2+^] Fig. 2C).

**Figure 2.**
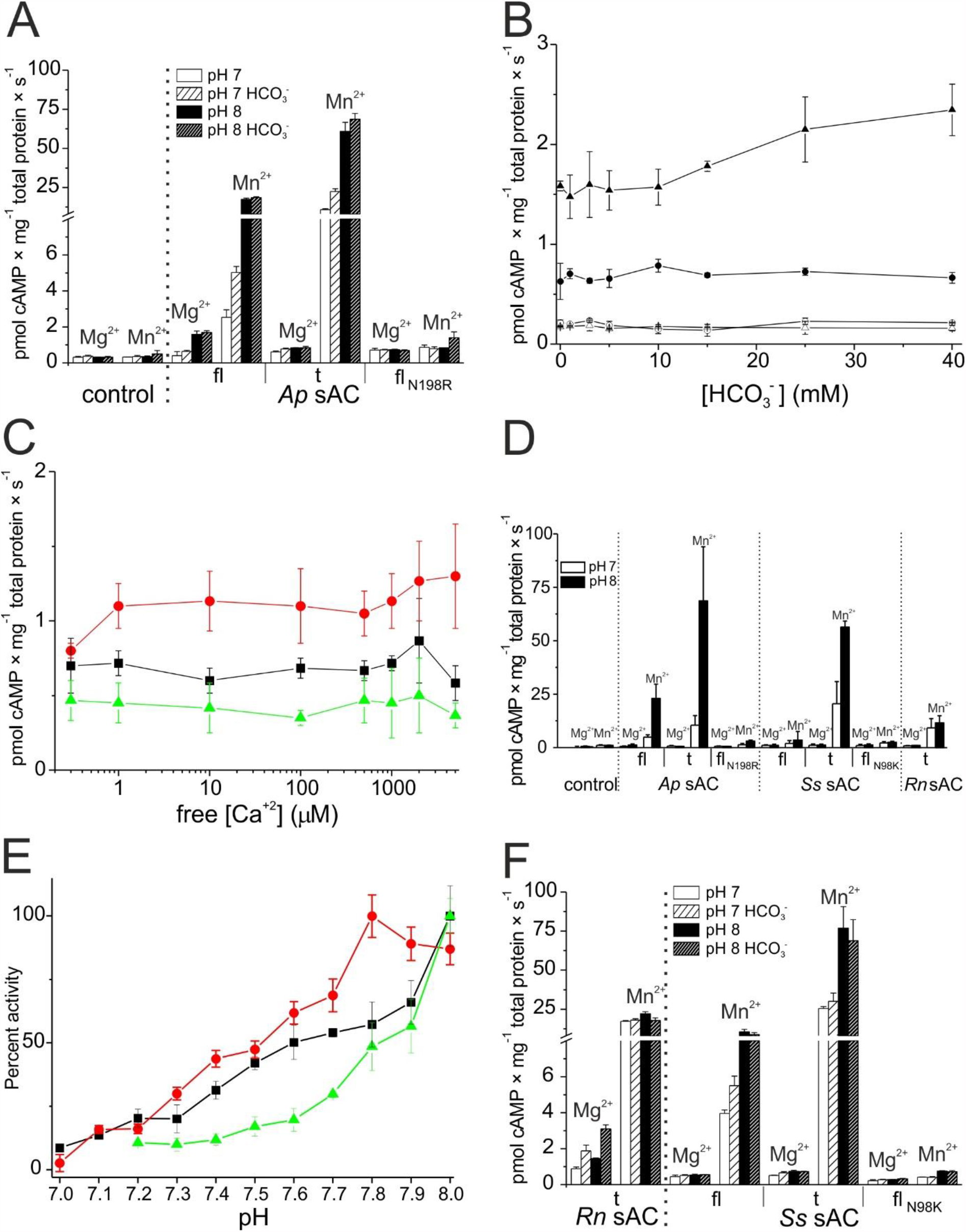
Heterologously expressed non-mammalian sAC orthologs are pH-sensitive and not activated by HCO_3_^-^ and Ca^2+^. **(A)** Bicarbonate sensitivity of heterologously expressed recombinant sAC_fl_, sAC_t_, and mutant N176R from *A. punctulata* at pH 7 and 8. MgATP (1 mM) or MnATP (1 mM) as substrate; HCO_3_^-^ 30 mM. **(B)** Bicarbonate dependence of cAMP synthesis of *Ap*sAC_fl_ at pH 7.1 (●) and 7.7 (▲). Control cells at pH 7.1 (○) and pH 7.7 (△). MgATP 1 mM. **(C)** Ca^2+^ sensitivity of sAC activity at pH = 8 of rat sAC_t_ (red), *Ap*sAC_fl_ (black), and control cells (green) in the presence of Mg^2+^ (5 mM) and ATP(2 mM). **(D)** Activity of sAC from *A. punctulata* (*Ap*sAC), *S. salar* (*Ss*sAC) and rat *Rn*sAC in lysates of HEK293 cells with MgATP (1 mM) and MnATP (1 mM) at pH 7 (white) and pH 8 (black), fl, full-length; t, truncated; fl_N198R_, full-length N198R mutant; fl_N98K_, full-length N98K mutant. **(E)** pH dependence of sAC activity in *Ap*sAC_fl_-transfected HEK cells (black), purified *Ap*sAC_t_ protein (red), and *A. punctulata* sperm (green). **(F)** Bicarbonate sensitivity of heterologously expressed recombinant sAC_fl_, sAC_t_, and mutant N98K from the salmon *S. salar* at pH 7 and 8. MgATP (1 mM) or MnATP (1 mM) as substrate.

We presumed instead that alkalization, a key step in chemotactic signaling of *A*.*punctulata* sperm (10, 23, 24), regulates sAC activity. Indeed, both truncated and full-length *Ap*sAC isoforms were exquisitely pH-sensitive (Fig. 2D); cAMP levels rose almost 10-fold when increasing the pH from 7 to 8 (Fig. 2E). In the freshwater spawner salmon *Salmo salar*, the *Ss*sAC regulation was similar: alkaline pH but not HCO_3_^-^ stimulated both *Ss*sAC_fl_ and *Ss*sAC_t_ activities (Fig. 2F).

In conclusion, the sAC from *A. punctulata* and *S. salar* is regulated by pH rather than HCO_3_^-^. Of note, other characteristic features such as membrane association, Mn^2+^ sensitivity, and higher catalytic activity of the truncated versus full-length form are shared by pH- and HCO_3_^-^- sensitive sACs (Fig. 2; Supplementary Fig. 1).

Residues R176 and K95 endow human sAC with HCO_3_^-^ sensitivity and are also involved in catalytic activity; replacement of both residues by alanine almost completely abolishes sAC activity (15). We tested whether HCO_3_^-^ sensitivity of *A. punctulata* and *S. salar* sACs can be recovered by replacing N198 (the R176 position) with Arg in *Ap*sAC_fl_ or N98 (the K95 position) with Lys in *Ss*sAC_fl_ (Fig. 2A, F). Both *Ap*sAC and *Ss*sAC mutants lost their pH regulation and Mn^2+^ sensitivity, but because they also lost most of their catalytic activity, it is unclear whether HCO_3_^-^ regulation was restored. In conclusion, the two residues are key to sAC activation in pH- and HCO_3_^-^-regulated sACs.

### A new cross-species sAC inhibitor

Small-molecule screens identified the mammalian sAC inhibitor LRE1 (17, 25). We found that the LRE1 congener TDI-8164 (Fig. 3A) inhibited pH-sensitive *Ap*sAC_t_ as well as HCO_3_^-^-sensitive *Rn*sAC_t_ and human *Hs*sAC_t_ with similar potency: the IC_50_ of TDI-8164 to inhibit *Ap*sAC_t_, *Rn*sAC_t_, and human *Hs*sAC_t_ was 0.6 ± 0.1 µM, 1.6 ± 0.36 µM, and 0.95 ± 0.1 µM, respectively (mean ± s.d., n = 3) (Fig. 3C-E). Of note, in intact cells, the IC_50_ to inhibit *Rn*sAC_t_ was 0.51 ± 0.08 µM (Fig. 3F), demonstrating that TDI-8164 readily permeates cell membranes.

**Figure 3.**
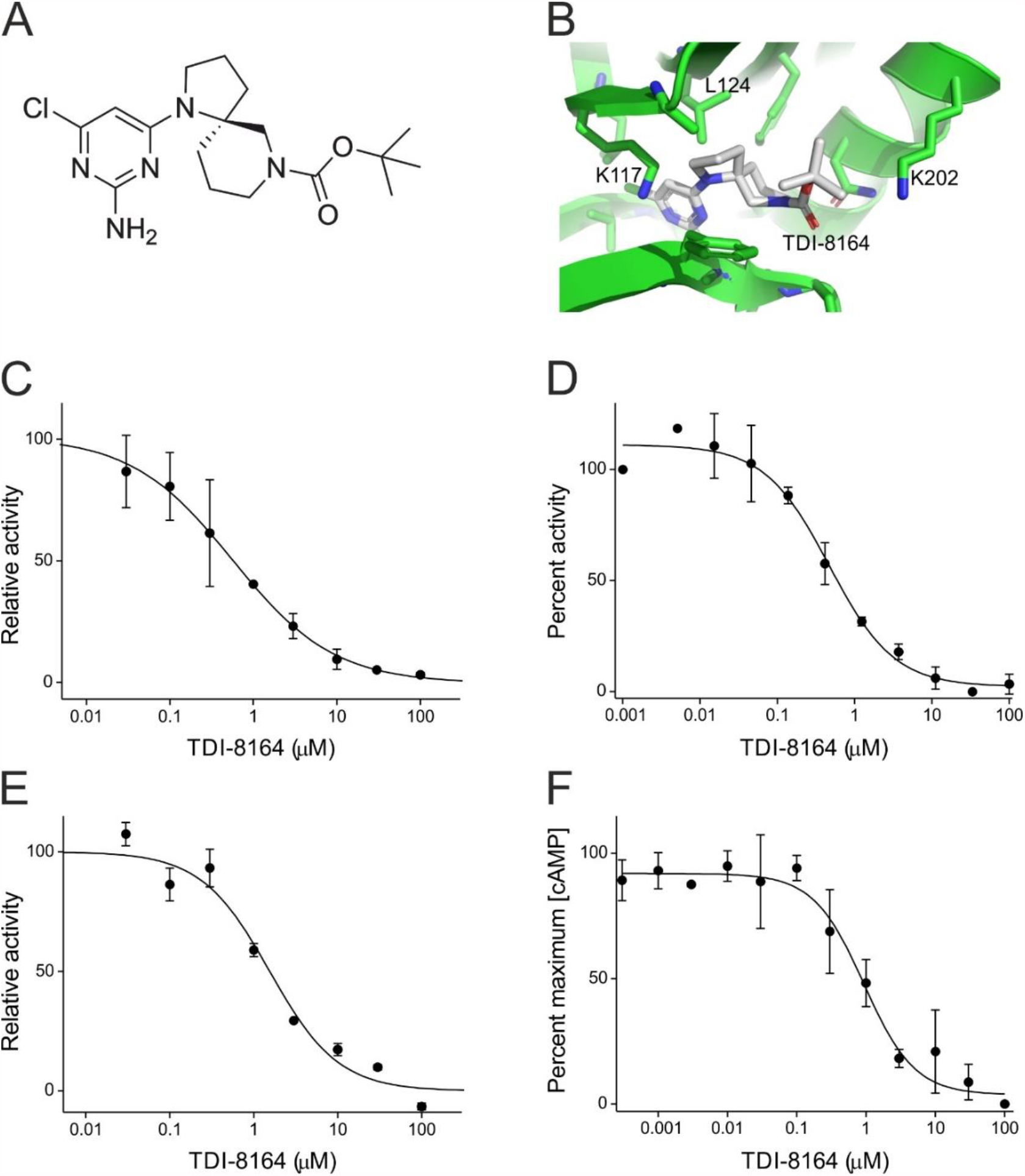
The drug TDI-8164 inhibits rat, human, and sea urchin sAC. **(A)** Chemical structure of TDI-8164. **(B)** Model of the *Ap*sAC catalytic core in complex with the inhibitor TDI-8164 (grey). Residues surrounding the compound are shown as sticks, and residues mentioned in the text are labeled. **(C-F)** Determination of IC_50_ value of cAMP synthesis for sAC inhibitor TDI-8164 using recombinant sAC from (C) sea urchin, (D) human, (E) rat in lysed HEK293 cells, and (F) rat in intact HEK293 cells. Mn^2+^ (5 mM), ATP (1 mM), pH 7.7. The IC_50_ = 0.6 ± 0.10 µM for *Ap*sAC in lysed HEK293 cells, 1.6 ± 0.36 µM for *Rn*sAC in lysed HEK293 cells, 0.51 ± 0.08 µM for purified *Hs*sAC protein, and 0.95 ± 0.1 for *Rn*sAC in intact HEK293 cells. Mean ± s.d. (n = 3) for panels C, E and n = 1-3 for panels D, F. Solid lines represent fit of a simple binding isotherm to the data.

Although two key residues in the HCO_3_^-^-binding site are altered in *Ap*sAC, the overall structure of the site remains mostly conserved (Fig. 1). Indeed, TDI-8164 fits well in the *Ap*sAC regulatory site based on a sAC-cat/LRE1 crystal structure (17) (Fig. 3B). The shared 2-amino-4-chloro moiety occupies the mammalian sAC bicarbonate site and interacts with conserved residues such as *Ap*sAC-K117 and L124, and the five-membered ring of TDI-8164 overlays with the LRE1 cyclopropyl group. The largest difference between the compounds is where the TDI-8164 six-membered ring and iso-butylester partly occupies the pocket accommodating LRE1’s thiophene group, but then extends further toward the active site. Contacts here include residues differing from mammalian sAC, such as *Ap*sAC-K202 (replacing N180), which likely explains why TDI-8164 showed high potency toward *Ap*sAC (Fig. 3B). In conclusion, TDI-8164 allows studying *Ap*sAC and other non-mammalian sAC orthologs in intact cells of species for which gene targeting is not available.

### cAMP synthesis in *A. punctulata* sperm is controlled by pH but not HCO_3_^-^

Considering that native sAC may adopt properties different from those *in vitro*, we examined the regulation of cAMP synthesis in intact motile *A. punctulata* sperm. Of note, in sea urchin sperm, tmAC isoforms are neither detectable at the protein level (9) see however (7, 13) nor functional level (Supplementary Information and Supplementary Fig. 2), whereas sAC is highly abundant (cytosolic concentration = 6.7 µM) (9). Consequently, cAMP synthesis in *A. punctulata* sperm is predominantly, if not exclusively, mediated by sAC (7, 26) simplifying its functional characterization in sperm.

We used quenched-flow technique (27, 28) to determine the change in cAMP content of sperm evoked by rapid mixing with artificial sea water (ASW, pH 7.8) containing HCO_3_^-^ (30 mM) or the chemoattractant peptide *resact*. Mixing with HCO_3_^-^ did not elevate cAMP levels, not even in sperm that had been preincubated with IBMX, a broadly-specific inhibitor of phosphodiesterases (PDE) that prevents breakdown of cAMP (Fig. 4A). By contrast, mixing with resact resulted in a transient cAMP increase (28, 29) that was greatly augmented by IBMX (Fig. 4A, B).

**Figure 4.**
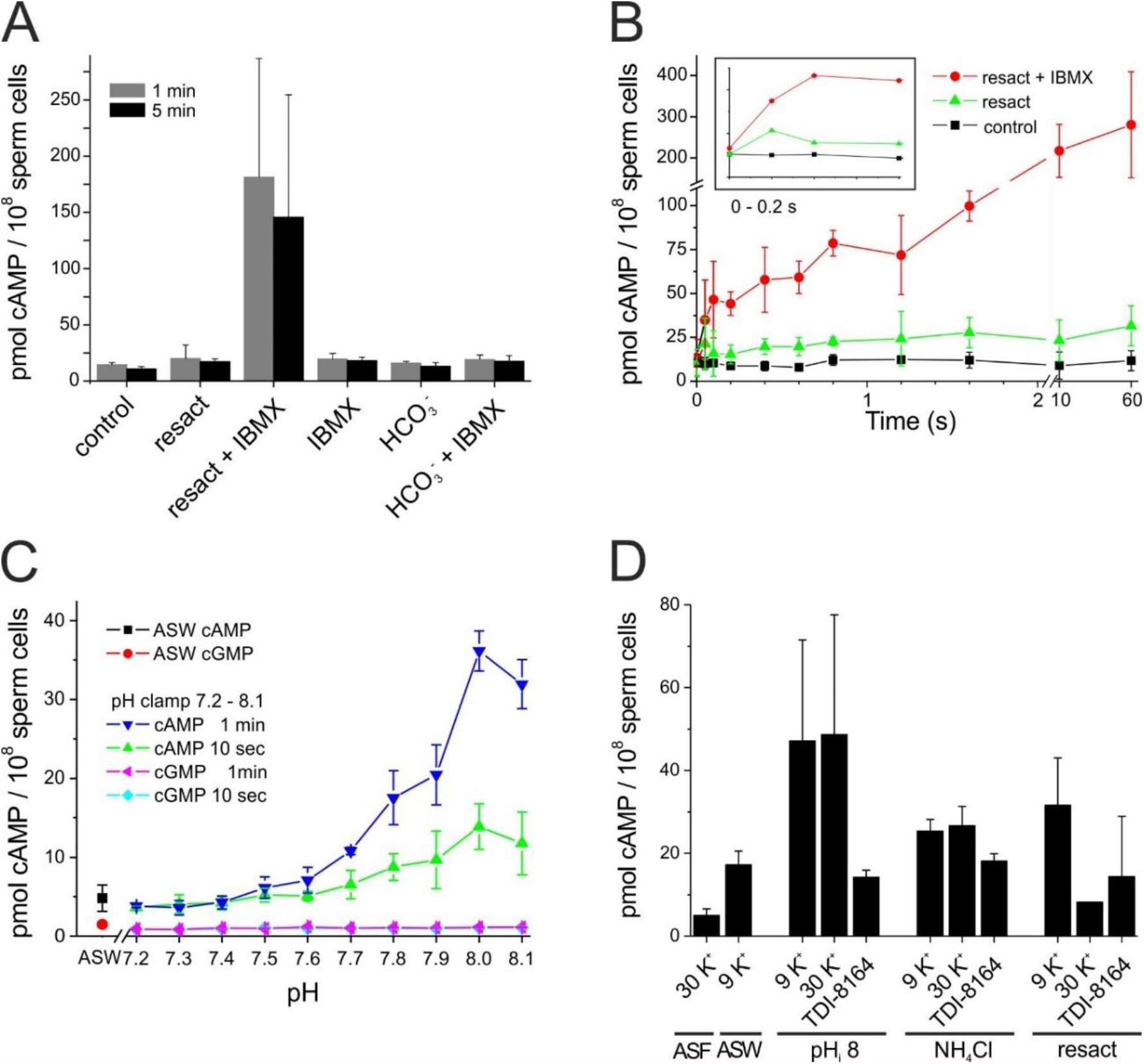
Control of cAMP in *A. punctulata* sperm. **(A)** The cAMP concentrations of unstimulated control sperm and after stimulation with HCO_3_^-^ (30 mM), HCO_3_^-^ (30 mM) + IBMX (1 mM), resact (250 nM), resact + IBMX (1 mM), and IBMX alone (1 mM) after 1 min (grey) and 5 min (black). **(B)** Time course of the changes in cAMP after rapid mixing with ASW (black), resact (250 nM, green), and resact + IBMX (1 mM) (red). Inset: blow-up of the first 200 ms; n = 3. **(C)** Control of cAMP and cGMP concentrations by intracellular pH_i_. The pH_i_ was adjusted by the pH_i_-clamp method. Changes in cAMP after 10 s (green) and 1 min (blue); changes in cGMP after 10 s (light blue) and 1 min (magenta). The control values in ASW are indicated as black (cAMP) and red (cGMP) dots; n = 3. **(D)** cAMP synthesis stimulated by pH_i_, NH_4_Cl, or resact and inhibited by high [K^+^] and the sAC inhibitor TDI-8164; n = 3, except n = 1 for resact + 30 mM K^+^. All data are the mean ± s.d. and the indicated number n of experiments.

Resact evokes a rapid hyperpolarization (30, 31) that alkalizes sperm via activation of a voltage-gated Na^+^/H^+^ exchanger (sNHE) (10, 24, 32). The resact-evoked hyperpolarization, alkalization, and rise of cAMP are abolished by high external [K^+^]_o_ (30 mM) (23, 24, 29, 33), suggesting that cAMP synthesis is stimulated by the change in voltage and/or pH_i_. Because sAC is tethered to the membrane, it may sense voltage changes. To distinguish between voltage and pH_i_ control, we studied the activity of sAC upon an increase in pH_i_ established by two different procedures. A rapid shift of pH_i_ from 7.2 to 8.1 was imposed onto sperm (“pH-clamp”) (24) using the pH_i_ pseudo-null-point method (34) (Supplementary Figs. 3, 4). Alternatively, sperm were alkalized by rapid mixing with the weak base NH_4_Cl. Both procedures elevated cAMP levels (Fig. 4C, D). The sAC inhibitor TDI-8164 suppressed both the alkaline- and the resact-induced cAMP increases (Fig. 4D), indicating that it is mediated by sAC; resact-induced cAMP synthesis was also inhibited by 30 mM K^+^, which inhibits hyperpolarization and, thereby, alkalization. To conclude, the activity of recombinant and native sAC is directly activated by alkaline pH_i_. Importantly, considering that the hyperpolarization causes alkalization, this result also unveils the long-standing mystery of voltage-sensitive cAMP synthesis in sperm (35).

The discovery that pH directly controls sAC provided a new starting point to study the role of cAMP in sperm of external fertilizers during spawning and chemotaxis.

### Sequence of signaling events during spawning

Non-motile sperm in seminal fluid become motile after spawning (36, 37). Phosphorylation of axonemal proteins by cAMP-stimulated PKA or activation of motor proteins by alkaline pH_i_ have been proposed to activate motility (38, 39). Considering that pH_i_ and cAMP are linked via the pH-sensitive sAC, we reasoned that the transition from the seminal fluid (SF) to sea water alkalizes sperm and raises cAMP levels. The ion concentrations of SF from *A. punctulata* (Supplementary Table 3) or other sea urchin species (40) are almost identical to that of sea water, except for K^+^ and pH. The [K^+^]_SF_ was 25.7 ± 2.1 mM (n = 5) vs. 9 mM in sea water, i.e., ASW. The pH of undiluted freshly spawned semen (i.e., sperm + seminal fluid) and seminal fluid (pH_SF_), was 6.8 ± 0.2 (n = 5) vs. 7.8 of ASW. We determined the sperm pH_i_ in artificial seminal fluid (ASF, which is ASW containing 27 mM K^+^ at pH 6.7) using the pH-clamp technique. The mean pH_i_ of sperm diluted in ASF and ASW was 6.84 ± 0.02 (n = 3) (Supplementary Fig. 4) and 7.2 (Supplementary Fig. 5) (24), respectively. Thus, sperm alkalize after spawning.

**Figure 5.**
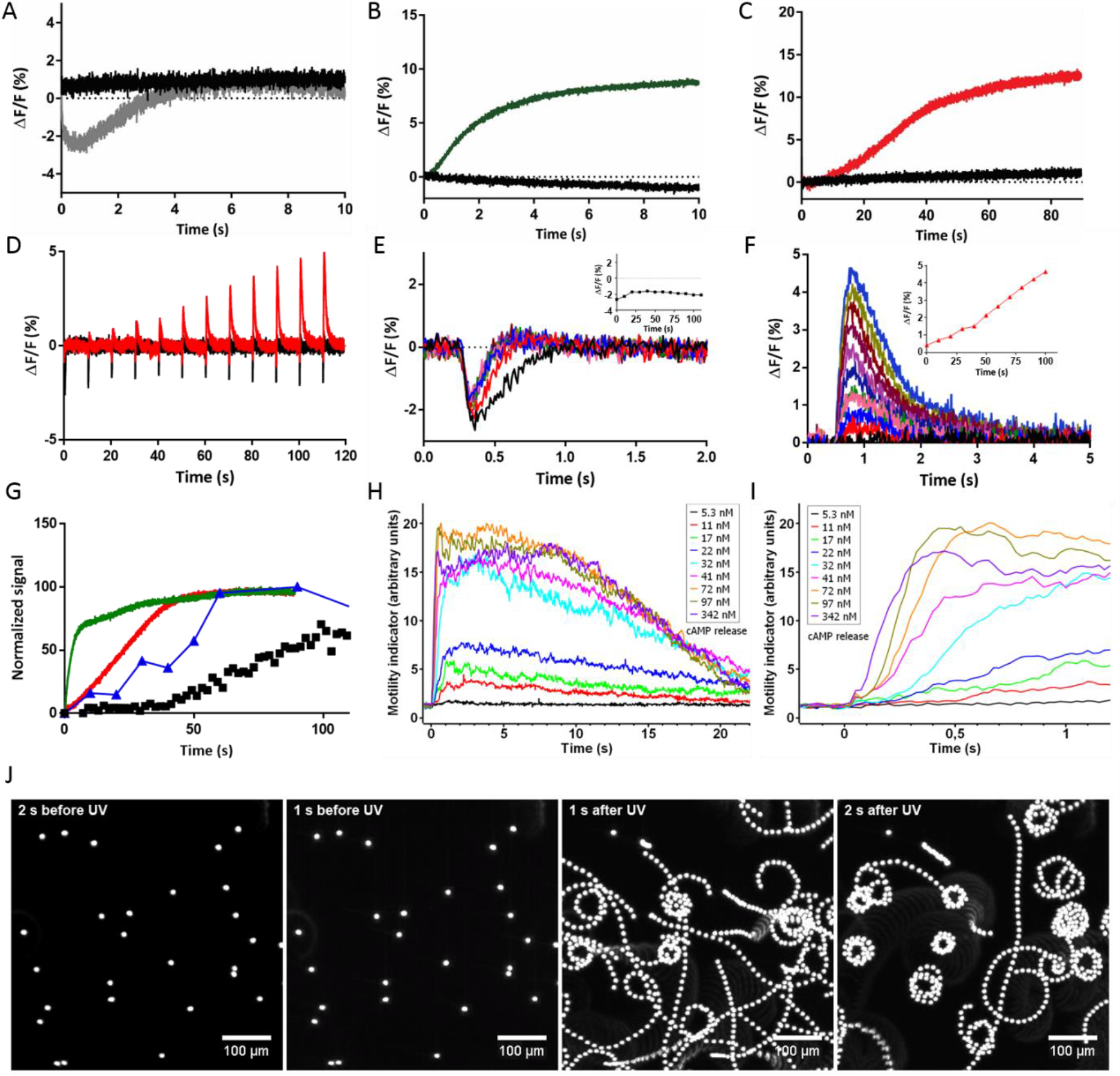
Cellular signaling events during spawning and activation of sperm. **(A)** Changes in V_m_ measured with the potentiometric probe FluoVolt after 1:2 mixing of sperm in ASF with 0KASW (gray) and 30 KASW (black). **(B)** Changes in pH_i_ measured with pHrodo Red after 1:2 mixing of sperm in ASF with 0KASW (red) or 30KASW (black). For comparison with Ca^2+^ signals, the pH_i_ signal has been inverted. **(C)** Changes in [Ca^2+^]_i_ measured with Fluo-4 after 1:2 mixing of sperm in ASF with 0KASW (red) or 30KASW (black). **(D)** cGMP-evoked V_m_ (black) and Ca^2+^ responses (red) after mixing sperm in ASW with 0KASW. **(E)** Superposition of cGMP-induced V_m_ responses evoked after 1:2 mixing of sperm in ASF with 0KASW; flash was applied in 10-s increments from t = 300 ms (black) to t = 110 s (turquoise). Inset: amplitude of V_m_ versus time. **(F)** Superposition of cGMP-induced Ca^2+^ responses evoked by the indicated times after 1:2 mixing of sperm in ASF with 0KASW color-coded as in panel E. Inset: amplitude of Ca^2+^ responses versus time. **(G)** Comparison of the time course of pH_i_ response (green), Ca^2+^ response (red), cAMP increase (blue), and motility (black) after dilution by ASW of sperm incubated in ASF. **(H)** Initiation of motility by UV light flashes of different energy in salmon sperm bathed in salmon ASF and loaded with DEACM-caged cAMP (20 µM), which raised cAMP concentrations from 5.3 nM to 342 nM. Each data point represents the mean from three animals and 6-9 experiments. **(I)** Extended timescale that illustrates the dose-dependent latency of the motor response. UV flash was delivered at t = 0. **(J)** Trajectories of salmon sperm from Supplementary Video 1 recorded 1 s and 2 s before (left) and after (right) release of cAMP by light.

To reveal the sequence of cellular events and the underlying mechanisms, we emulated spawning of *A. punctulata* sperm using rapid mixing techniques (27), while following the changes in V_m_ and pH_i_ using fluorescent indicators. Before mixing, sperm in ASF were loaded with the respective fluorescent probes and then diluted with ASF. The dilute sperm suspension was rapidly mixed 1:2 in the stopped-flow device with K^+^-free ASW (0KASW, pH 8.0); thus, upon mixing, [K^+^] dropped to 10 mM. As control, sperm were mixed with ASW containing 30 mM K^+^ (30KASW, pH 8.0), leaving [K^+^] unchanged. For both mixing protocols, pH_o_ was stepped from 6.7 to 7.7.

The [K^+^]_o_ controls the resting V_m_ of sea urchin sperm (31); therefore, sperm are predicted to hyperpolarize during spawning due to the lower [K^+^] of ASW. Indeed, mixing with 0KASW, evoked a rapid transient hyperpolarization (Fig. 5A). The hyperpolarization was accompanied by a pronounced intracellular alkalization (Fig. 5B). Both, hyperpolarization and alkalization were completely abolished by mixing with 30KASW (Figs. 5A, B). This observation is remarkable as it shows that protons do not passively redistribute across the sperm membrane by conventional Na^+^/H^+^ exchange. Instead, a change in V_m_ is required, which activates the voltage-gated Na^+^/H^+^ exchanger SLC9C1 (9, 10). Next, we examined whether the changes in V_m_ and pH_i_ during spawning affect the voltage- and pH-gated Ca^2+^ channel CatSper (24). On mixing with ASW, after some latency, [Ca^2+^]_i_ rose continuously and reached a plateau after 90 s (Fig. 5C). Finally, we followed the changes in cAMP level after mixing with ASW. The resting cAMP level of sperm in ASF was extremely low (3.3 ± 1.5 pmol/10^8^ cells, n = 2) and rose about 5-fold (15.1 ± 4.0 pmol/10^8^ cells) with a half-time of approximately 50 s after dilution into ASW (Fig. 5G). In conclusion, spawning first triggers a change in V_m_, followed by a rise of pH_i_ and finally a stark rise of cAMP and [Ca^2+^]_i_.

### Sperm motility is initiated by alkalization and a rise of cAMP

Next, we examined how changes in extracellular [K^+^] and pH_i_ during spawning affect motility. Upon dilution of dry sperm into ASW (1:200), 87.5 ± 5.3% (n = 3, 740 cells) of sperm became motile, whereas dilution into ASF did not activate motility, demonstrating that the ion composition of SF rather than an unknown inhibitory factor keeps sperm quiescent. Motility activation was not instantaneous but started after about 40 s (Fig. 5G). After dilution into ASF at pH 7.8, or in ASW at pH 6.7, or in ASW containing 30 mM K^+^ (30KASW) only about 4-15% of sperm became motile (ASW 6.7: 9.9 ± 4.1%; 30 KASW: 17.6 ± 14.8%; (n = 3) 490-536 cells). These results show that the pH_o_ difference between SF and ASW is not sufficient to initiate sperm motility and a decrease of [K^+^]_o_ is required. Thus, sperm motility and cAMP synthesis by sAC are triggered by the same mechanisms.

We tested whether initiation of motility also requires cAMP in salmon sperm, which become motile upon dilution into freshwater. Immotile salmon sperm bathed in salmon ASF and loaded with DEACM-caged cAMP became motile in a dose-dependent fashion upon UV-uncaging cAMP (Fig. 5H, I, Supplementary Movie 1). At high cAMP concentrations, the latency of the motor response was as short as 100 ms. Of note, because a pH_i_ change was bypassed, a cAMP rise is sufficient to initiate motility. After the cAMP rise, sperm swam for about 20 s on curvilinear trajectories (Fig. 5J) until they became immotile again (Fig. 5H), reminiscent of the transient spawning-evoked motility of sperm from several other fish species (36). Our results reveal that external fertilizers share a common mechanism of motility initiation that involves a pH-induced rise of cAMP.

### The chemotactic signaling pathway matures after spawning

The relatively slow cellular events after spawning (Figs. 5A-C) may represent a maturation process reminiscent of capacitation of mammalian sperm inside the female reproductive tract (2). We probed the chemotactic cGMP-signaling pathway (9) in time using caged cGMP as a photo-trigger. Two key events - hyperpolarization and Ca^2+^ response - were probed by brief light flashes that released intracellular cGMP in sperm loaded with caged cGMP. Flashes were delivered 300 ms to 120 s after dilution into ASW (Fig. 5D). This stimulation protocol monitored the recruitment of signaling components. The amplitude and time course of the cGMP-evoked V_m_ responses did not change with time after dilution, except that V_m_ responses evoked during the first 10 s recovered more slowly (Fig. 5E). Thus, the cGMP-gated CNGK channel is almost instantaneously functional after dilution/spawning. By contrast, at times ≲20 s after mixing, cGMP evoked no or very small Ca^2+^ responses whose amplitude grew with time and reached a maximum after about 90 s (Fig. 5D, F). The slow increase of basal [Ca^2+^]_i_ after mixing matched the increase of the cGMP-evoked Ca^2+^ responses (Fig. 5C, F). In conclusion, after spawning, CatSper channels are quiescent and require some time to become functional. The exquisite pH sensitivity of sea urchin CatSper (24) suggests that spawning shifts pH_i_ to a regime where voltage-dependent activation of CatSper channels is enabled. However, the rise of pH_i_ after mixing is almost complete within 10 s, whereas full recruitment of functional CatSper channels requires approximately 90 s (Fig. 5G). Therefore, mechanisms in addition to a pH_i_ shift may contribute to full CatSper functionality.

### cAMP promotes full recovery from hyperpolarization and the Ca_2+_ response

The hyperpolarization-activated and cyclic nucleotide-gated (HCN) channel carries an inward Na^+^ current that ultimately terminates chemoattractant-induced hyperpolarization (9, 12). During recovery from hyperpolarization, CatSper channels open and Ca^2+^ flows into the flagellum (24, 31). While hyperpolarization activates HCN channels, cAMP greatly enhances the open probability (12). We examined whether cAMP synthesis by sAC is required for HCN function by recording V_m_ responses in the presence of the sAC inhibitor TDI-8164. The compound did not affect the kinetics of the resact-induced hyperpolarization, showing that cAMP is not required for activation of the pathway (Fig. 6A). By contrast, recovery from hyperpolarization was incomplete and markedly delayed by TDI-8164 (Fig. 6A). At stronger stimulation, recovery accelerated; the acceleration was prevented in the presence of TDI-8164 (Fig. 6B). Slow and incomplete recovery should affect the Ca^2+^ response. Indeed, at low resact concentrations, the resact-evoked Ca^2+^ response is abolished by TDI-8164, and at high resact concentrations, the Ca^2+^ response is drastically delayed and sluggish (Fig. 6C, D). As a control, TDI-8164 did not inhibit alkalization that is important for CatSper activation (Fig. 6E, F); instead, the pH_i_ signals in the presence of TDI-8164 were slightly larger than those of the control (Fig. 6E, F). In conclusion, cAMP is required for quick and complete V_m_ recovery and, thereby, shapes the chemotactic Ca^2+^ response.

**Figure 6.**
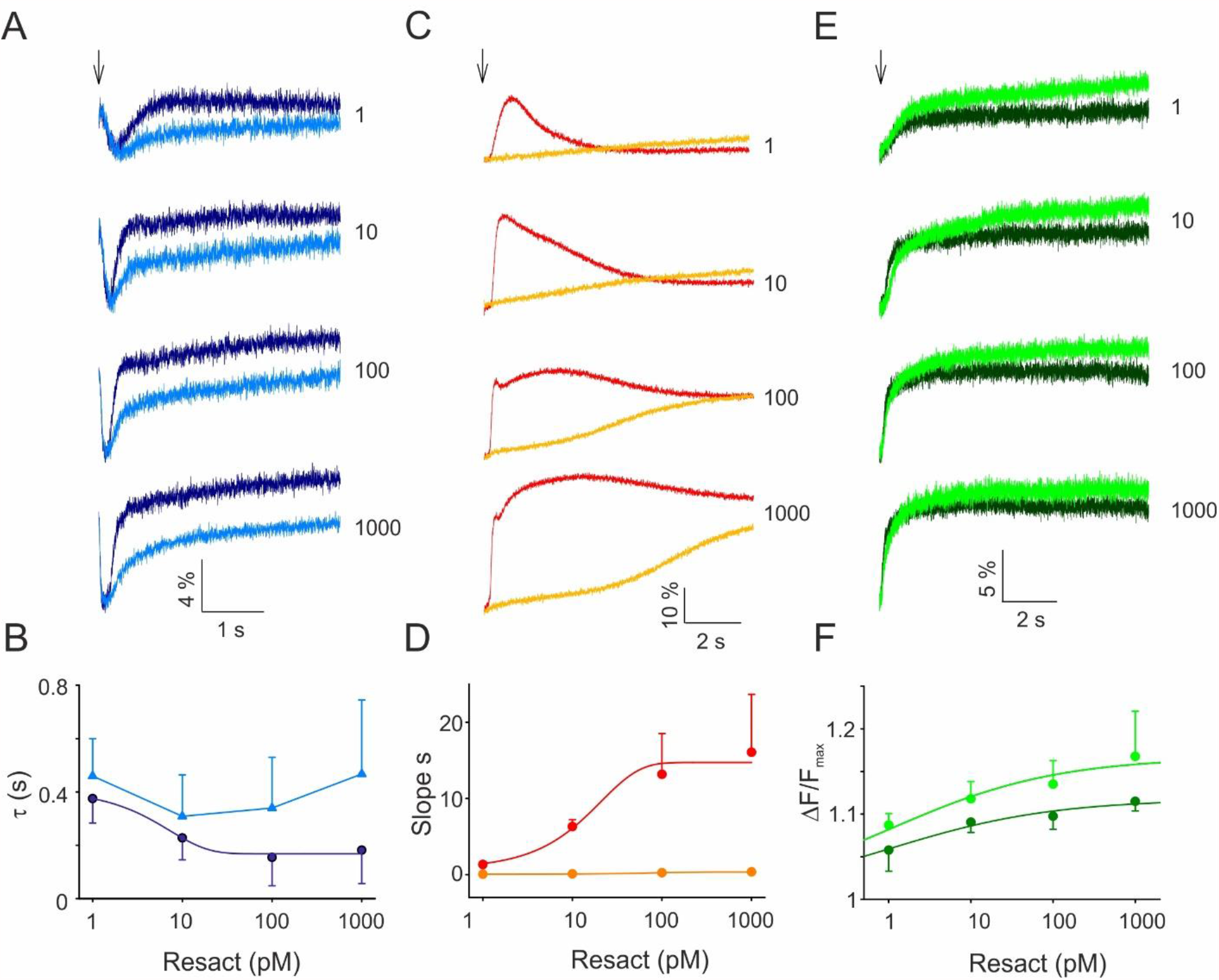
The sAC inhibitor TDI-8164 slows V_m_ recovery and impairs Ca^2+^ response. **(A)** Resact-evoked hyperpolarization in the absence (dark blue) and presence (light blue) of TDI-8164 (30 μM); resact concentrations are indicated in pM; voltage-sensitive dye Di-8-ANEPPS (2 μM). **(B)** Time constant of repolarization in control (dark blue) and TDI-8164-treated (light blue) sperm. **(C)** Resact-evoked Ca^2+^ responses in control (red) and TDI-8164-treated sperm (orange); resact concentrations as indicated; Ca^2+^ dye Fluo-4 (10 μM). **(D)** Initial rate of Ca^2+^ increase in control (red) and TDI-8164-treated sperm (orange) in dependence on resact concentration. **(E)** Resact-evoked alkalization in the absence (dark green) and presence (light green) of TDI-8164 (30 μM); BCECF dye (10 μM). **(F)** Relative changes ΔF/F_max_ of BCECF without (dark green) and with TDI-8164 (light green). All data are mean ± s.d. (n = 3)

## Discussion

Here, we show that metazoan sAC orthologs fall into two functionally distinct subfamilies. The original sAC family first described in mammalian sperm is directly activated by HCO_3_^-^ and is not pH sensitive. The new sAC subfamily identified here is activated by alkaline pH and is not sensitive to HCO_3_^-^. Thus, the pH-sensitive sAC serves as a direct pH_i_ sensor. In the pH-sensitive sAC, either K95 or R176, which are key for HCO_3_^-^ binding and activation of mammalian sAC, is exchanged for an asparagine residue, suggesting that these substitutions are responsible for the loss of HCO_3_^-^ sensitivity and the switch to pH sensitivity. The species across ten phyla using the pH_i_-regulated sAC comprise a large number of external fertilizers, suggesting that internal fertilizers such as mammals employ HCO_3_^-^-sensitive sAC, whereas external fertilizers that spawn into aquatic environment low in HCO_3_^-^ carry the pH-sensitive sAC. There are notable exceptions: reptiles fertilize internally; accordingly, some species such as lizards and snakes feature the R176/K95 pair, which predicts HCO_3_^-^ sensitivity, whereas in crocodiles this amino-acid pair is not conserved. It will be interesting to examine whether the sAC from either reptiles fall along these lines or whether fertilization mechanisms are different.

Apart from coordinating HCO_3_^-^, the R176/K95 pair is also required for catalytic function. In the human K95A/R176A mutant, the HCO_3_^-^ affinity is lowered only by 2-3-fold, yet the catalytic activity is almost completely abolished (15). In the pH-sensitive sAC, the substituted residues at these sites contribute to pH_i_ sensitivity. Substitutions to the mammalian genotype – N98K (salmon) and N198R (sea urchin) – although unsuccessful at restoring HCO_3_^-^ regulation, did abolish pH sensitivity and catalytic activity in general. We speculate that these two residues act in concert with additional regions in the polypeptide to confer either HCO_3_^-^ or pH sensitivity. However, the overall structure of the regulatory HCO_3_^-^/H^+^ site is maintained as TDI-8164, which targets this site, inhibits both HCO_3_^-^- and pH-sensitive sAC with similar potency.

The evolution of sAC is intriguing. The sAC of cyanobacteria carries K95 and R176 and is regulated by HCO_3_^-^ and Ca^2+^ (19, 41, 42). Cyanobacteria rely on CO_2_ and HCO_3_^-^ as carbon source for photosynthesis (43). They evolved several CO_2_-concentrating mechanisms that improve photosynthetic performance and were essential for survival when CO_2_ levels markedly dropped around 350 Mya (44). The products of the *SbtAB* operon link cAMP with carbon import: association of cAMP-liganded SbtB with the Na^+^/HCO_3_^-^ cotransporter SbtA inhibits HCO_3_^-^ uptake (43, 45). When the primordial sAC was adopted by metazoan for fertilization in the sea, with a few exceptions, either one of these two amino acid sites has been changed to allow for pH-rather than HCO_3_^-^control. Finally, the two amino-acid substitutions were restored during evolution of mammals along with bicarbonate sensitivity. Thus, the HCO_3_^-^-sensitive sAC probably represents an adaptation to internal fertilization in the high-HCO_3_^-^ environment of the female reproductive organ.

The function and regulation of cAMP in non-mammalian sperm have been elusive. Here we identify several different functions along the path to fertilization (Fig. 7). First, cAMP is very low in quiescent sperm and rises after spawning. A rise of cAMP or pH_i_ has been considered necessary for sperm to become motile (38, 46, 47). The discovery of a pH-sensitive sAC reconciles these seemingly contradictory mechanisms. The pH_i_ rise happens ahead of the cAMP increase and motility activation, arguing that alkaline-stimulated cAMP synthesis followed by PKA-mediated phosphorylation of dyneins or associated proteins (38, 48) initiates motility. In support of this notion, a flash-induced rise of cAMP alone almost instantaneously activated vigorous motility in salmon sperm.

**Figure 7.**
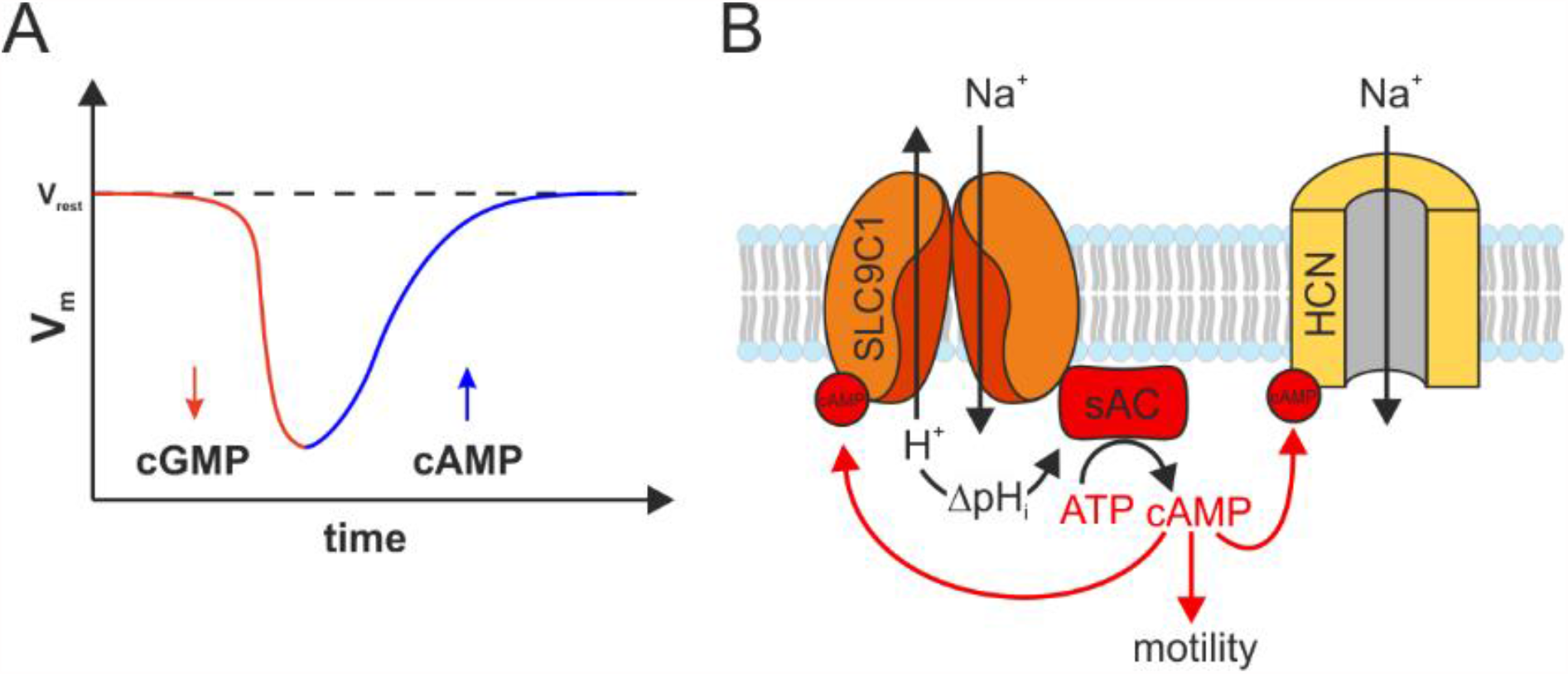
Model of sAC function and cAMP targets. **(A)** The hyperpolarizing V_m_ response is shaped by two ion channels and two cell messengers: cGMP opens CNGK channels that cause hyperpolarization, and cAMP opens HCN channels that cause depolarization. **(B)** The Na^+^/H^+^ exchanger SLC9C1 and the sAC reciprocally control each other’s activity. cAMP shifts V_1/2_ of sNHE activation, and sNHE controls sAC via pH_i_. It enhances the open probability of HCN channels and, thereby, contributes to CatSper channel opening during recovery from chemoattractant-evoked hyperpolarization; finally, cAMP controls dynein motor proteins via phosphorylation.

Second, the spawning-induced cAMP rise may promote CatSper maturation. Both alkalization and a transient hyperpolarization are required to open CatSper channels (24). Alkalization is mediated by the sNHE exchanger, and hyperpolarization is terminated by hyperpolarization-activated and cyclic nucleotide-gated (HCN) channels (12, 49). Both, sNHE exchanger and HCN channels are activated by hyperpolarization, and their activation is directly modulated by binding of cAMP to a cyclic nucleotide-binding domain (10, 12). The slow rise of pH_i_ and cAMP after spawning shifts the voltage dependence of sNHE, HCN channels, and CatSper to the final permissive voltage range.

Third, cAMP is transiently elevated after chemoattractant stimulation (Fig. 4B) (28). The role of cAMP in the chemotactic signaling pathway has been elusive. Pharmacological inhibition of the HCN channel (9) or cAMP synthesis prevents or slows down the recovery from hyperpolarization and impairs the Ca^2+^ response (Fig. 6). Because activity of the heterologous HCN channel is greatly enhanced by cAMP (12), we conclude that the cAMP rise promotes recovery from hyperpolarization. Thus, electric signaling in sperm is a tug-of-war between two ion channels and two cellular messengers: cGMP opens K^+^ channels that hyperpolarize the cell, whereas cAMP opens HCN channels that blunt hyperpolarization (Fig. 7A).

Fourth, cAMP sets the voltage range of sNHE activity (10). Now, we identify the pH-sensitive sAC as a new target of Na^+^/H^+^ exchange. Thus, sNHE and sAC reciprocally control each other (Fig. 7B). The reciprocal control of sNHE and sAC may be conserved across phyla. Sperm from sNHE^-/-^ mice lack the full-length sAC_fl_, and are immotile, rendering males infertile (50, 51). However, a rise of cAMP achieved by various techniques rescues motility of sNHE^-/-^ sperm and the infertility phenotype *in vitro* (50, 52, 53), suggesting a molecular and functional interaction between sAC and sNHE (50). However, because mammalian sAC is not pH-sensitive (42) and mammalian sNHE may not facilitate Na^+^/H^+^ exchange (10), reciprocal control must be mechanistically different.

On a final note, most external fertilizers living in an aquatic environment are directly exposed to changes in the CO_2_/HCO_3_^-^/H^+^ equilibrium. The increase of dissolved CO_2_ due to climate change can affect pH_i_ homeostasis in two ways. The reaction of CO_2_ with H_2_O to form HCO_3_^-^ and H^+^ acidifies the aquatic habitat. Moreover, CO_2_ readily crosses cell membranes and acidifies the cytosol. A permanent disturbance of cell pH_i_ is expected to affect sAC function and, thereby, reproduction of aquatic organisms (54).

## Author Contributions

H.G.K., H.H., and U.B.K. designed the project. H.H., U.B.K., R.S., and T.S. designed, performed, and analyzed quenched-and/or stopped-flow experiments. H.G.K. biochemically and functionally characterized the recombinant sACs. W.B. designed and cloned all sACs. L.A. and R.P. designed, performed, and analyzed the single-cell motility experiments. M.M., A.J. and A.S. were responsible for the identification, docking studies, and synthesis of TDI-8164. J.B. and L.R.L. developed and N.K. characterized the TDI-8164 inhibitor. C.S. performed the homology modeling. C.K. characterized the purified *Ap*sAC proteins. U.B.K. wrote the manuscript. All authors revised the manuscript for important intellectual content and approved the manuscript.

## Acknowledgments

We thank Heike Krause for preparing the manuscript and D. Fey and J. Hellmann (LANUV Kirchhundem-Albaum), and A. Nemitz (Wildlachszentrum Siegburg, Rheinischer Fischereiverband, Stiftung Wasserlauf, Wahnbachtalsperren-Verband, Wanderfischprogramm NRW) for providing salmon sperm. Financial support by the Deutsche Forschungsgemeinschaft *via* the priority program SPP1726 ‘‘Microswimmers’’ (to U.B.K.) and STE1701/11 (to C.S.), from the United States National Institutes of Health R01 HD088571 and P50 HD100549 (to J.B. and L.R.L), and financial support of TDI from Takeda Pharmaceutical Company, Memorial Sloan Kettering Cancer Center, the Rockefeller University, Weill Cornell Medicine, Mr. Lewis Sonders, and other philanthropic sources, is gratefully acknowledged.

## Methods

### Homology modeling of *Ap*sAC and its TDI-8164 complex

A homology model of *Ap*sAC was generated with Modeller v9 (55), using the closest homologs from the PDB as templates: CyaC from *S. platensis* (PDB ID 1WC0) (19) for C1 and human sAC-C2 (PDB ID 4CLL) (15) for C2. A model of an *Ap*sAC complex with the inhibitor TDI-8164 was generated by overlaying the *Ap*sAC1 model with a *hs*sAC-cat structure in complex with the related compound LRE1 (PDB ID 5IV4) (17). Coordinates and parameters for TDI-8164 were generated with ProDrg2 (56) and the compound was superposed on LRE1, followed by manual adjustment of the compound conformation within the binding site.

### Identification of TDI-8164 (tert-butyl (*R*)-1-(2-amino-6-chloropyrimidin-4-yl)-1,7-diazaspiro[4.5]decane-7-carboxylate)

TDI-8164 is the sole active enantiomer of a racemic mixture originally identified as a potent sAC inhibitor that was inert against mammalian tmACs (data not shown). The parent racemic compound was derived from a high-throughput organic synthesis (HTOS) of ∼50 LRE1 analogs. Briefly, a library of 1000 compounds was computationally enumerated based on the 2-amino-6-chloropyrimidine scaffold of LRE1 with 500 amine and 500 aldehyde building blocks provided by a CRO (Axcelead). The enumerated compounds were then computationally docked to the crystal structure of hsAC (PDB ID 5IV4) using rDock (http://rdock.sourceforge.net/), energy minimized, and re-scored by induced-fit docking energy in MOE (https://www.chemcomp.com/Products.htm). A diverse subset of 48 compounds was selected for synthesis from the top-scoring 100 compounds in each enumeration set, and evaluated by the RapidFire Mass Spec hsAC assay.

### Synthesis of TDI-8164 (tert-butyl (*R*)-1-(2-amino-6-chloropyrimidin-4-yl)-1,7-diazaspiro[4.5]decane-7-carboxylate)

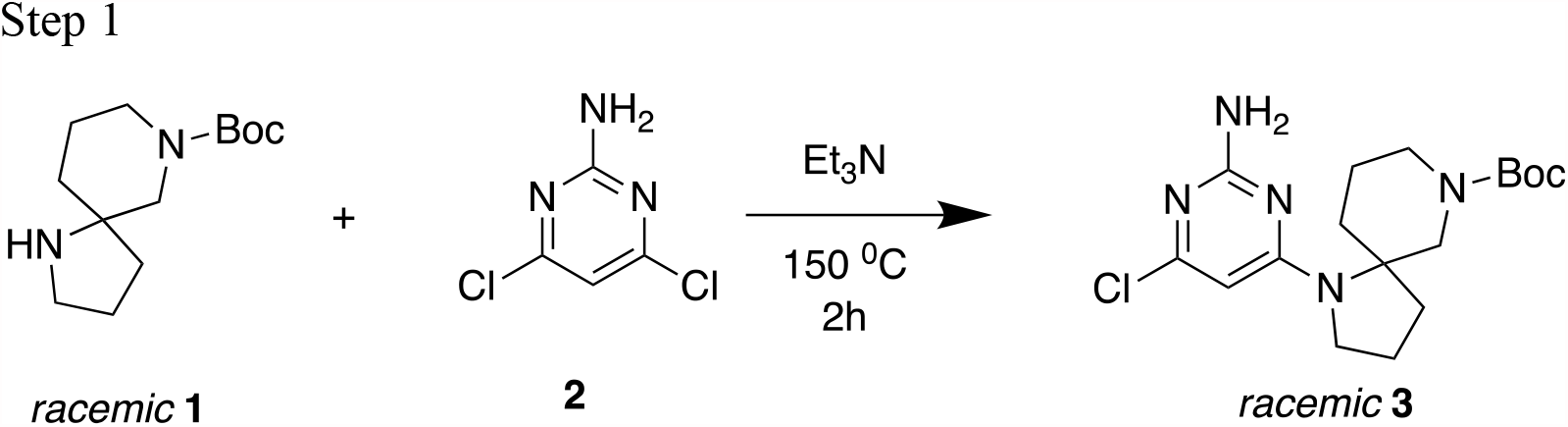

Compound **2** (1.02 g, 6.24 mmol, 3 *eq*), Et_3_N (630 mg, 6.2 mmol, 0.87 mL, 3 *eq*) and *racemic* **1** (500 mg, 2.08 mmol, 1 *eq*) were taken up into a microwave tube. The sealed tube was heated at 150°C for 2 h under microwave irradiation. The mixture was cooled to 25°C. The reaction was repeated four times, and the resulting crude mixtures were purified by preparative-HPLC (column: Kromasil Eternity XT 250, 80mm x 10 mm; mobile phase: [water (0.05% ammonia hydroxide v/v)-ACN];B%: 60%-80%, 33-minute gradient) to afford *racemic* **3** (2.30 g, 5.91 mmol) as a brown solid. LCMS (M+H)^+^ 368.3.

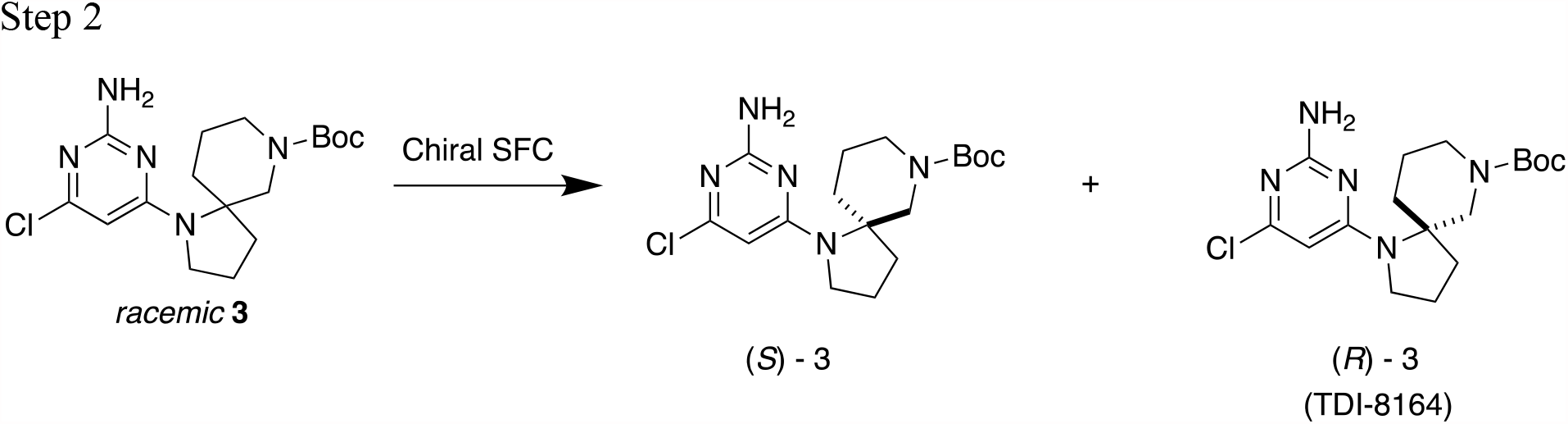

*Racemic* **3** (2.30 g, 5.91 mmol) was purified by chiral preparative SFC (column: DAICEL CHIRALPAK AD (250 mm x 30 mm, 10 mm); mobile phase: [0.1% NH_3_H_2_O in EtOH]; B%: 40%, isocratic elution) to afford (*S*)-**3** (Peak 1: 0.863 g, 2.30 mmol, 37 % yield) and (*R*)-**3** (Peak 2: TDI-8164, 0.899 g, 2.43 mmol, 39 % yield) as yellow solids. ^1^H NMR (400 MHz, CDCl_3_) δ: 5.77 (s, 1H), 4.76 (s, 2H), 4.17 - 4.03 (m, 2H), 3.74 - 3.72 (m, 1H), 3.46 - 3.37 (m, 2H), 3.22 - 3.21 (m, 1H), 2.69 (m, 1H), 2.21 - 2.18 (m, 1H), 1.92 (s, 2H), 1.75 - 1.69 (m, 2H), 1.63 (m, 1H), 1.46 (s, 9H), 1.41-1.38 (m, 1H).

### Cloning of full-length and truncated sAC and sAC mutants

Three sets of primer pairs, designed using genomic data of *A. punctulata*, were used to obtain the full-length sequence of soluble adenylate cyclase (sAC) by PCR amplification on a *A. punctulata* testis library. Primers C3299 and C3302 were used to amplify the 5′ part of the sequence (bp 1 to 1790). Primer C3299 introduces a BamHI site followed by a perfect Kozak sequence preceding the start codon. Primer C3302 introduces an EcoRI site by silent mutation. Primers C3307 and C3310 were used to amplify bases 1768 to 3717. Primer C3310 introduces a XbaI site by silent mutation. The 3′ part (bp 3704 to 5556) was amplified with primers C3315 and C3318, followed by a PCR with primers C3315 and C1001, which adds a sequence for an HA-tag to the 3’ end followed by a stop codon and a NotI site. The primer sequences were:

CACCGGATCCACCATGAGTGAGGCTATAAACTCAACACAG (C3299) ACCAGAATTCTCCGATGGTCTCCTCTCTCTCGC (C3302) GGAGACCATCGGAGAATTCTGGTGTTCCAGGG (C3307) GCCTTCTAGAGGTTCATCATCATCTGAGGAATC (C3310) AACCTCTAGAAGGCAGACCCAAGGAATATAAC (C3315) TAATAAGCGGCCGCTAGGCGTAGTCGGGCACGTCGTAGGGGTATTCCTCTTCACCCCTGGGGAGAG (C3318) TAATAAGCGGCCGCTAGGCGTAGTCGGGCACGTCGTAGGGG (C1001)

The three resulting PCR fragments were cloned into vector pcDNA6/V5-HisA (Invitrogen, Carlsbad, USA) digested with BamHI and NotI to obtain the full-length clone. The DNA for the *S. salar* sAC was synthesized according to the annotated sequence for the soluble adenylate cyclase 10, XM_014151728. A HA-tag was added to the C-terminal end. The DNA was cloned via HindIII and XbaI sites into vector pcDNA3.1/zeo(+) (Invitrogen, Carlsbad, USA). Mutations were introduced into the sequences by standard methods.

pc6-ApsACN198R was constructed using primers C3299, C4097 and C4098, which introduces the mutation N198R, and primer C1001. The resulting PCR fragment was cloned into vector pcDNA6/V5-HisA (Invitrogen, Carlsbad, USA). The primer sequences were:

CACCGGATCCACCATGAGTGAGGCTATAAACTCAACACAG (C3299) CAAACTTCTCGGCGATTCTAGCCTCCAGTACTG (C4097) CAGTACTGGAGGCTAGAATCGCCGAGAAGTTTG (C4098) TAATAAGCGGCCGCTAGGCGTAGTCGGGCACGTCGTAGGGG (C1001)

Truncated sAC from Arbacia punctulata (Met 1 to Leu 577) was constructed by successively PCRs using 5’ primer C3299 and 3’ primers C3680, C3681, C3682. Primer C3680 matches to the ApsAC up to L577 and adds a part of the HA-tag, primer C3681 adds part of the HA-tag, followed by a short linker (Gly, Ser, Gly), primer C3682 adds a hexa His-tag, followed by a stop codon and a XbaI site. The resulting PCR fragment was cloned into vector pcDNA6/V5-HisA (Invitrogen, Carlsbad, USA). The primer sequences were:

CACCGGATCCACCATGAGTGAGGCTATAAACTCAACACAG (C3299) GGGCACGTCGTAGGGGTACAAGAAGATAGACATCTCCTTGTCTCG (C3680) GGTGGCCGCTGCCGGCGTAGTCGGGCACGTCGTAGGGGTACAAG (C3681) ATATCTAGATTAGTGGTGGTGGTGGTGGTGGCCGCTGCCGGCGTAG (C3682)

pc3Z-SssACN98K was constructed using primers C3978, C4099, C4100 and primer C0609. Primer C3958 introduces a HindIII site followed by a perfect Kozak sequence preceding the start codon, Primers C4099 and C4100 introduce mutation N98K. Primer C0609 anneals to the HA tag. The resulting PCR fragment was cloned into vector pcDNA3.1/zeo(+) (Invitrogen, Carlsbad, USA). The primer sequences were:

TAAGCTTCCACCATGGGCTGGATCAAGGGCGACGGCGAGATCGAG (C3978) CGTCGCCGGCGTACTTCAGGATGTCGCC (C4099) GGCGACATCCTGAAGTACGCCGGCGACG (C4100) TCTTCTAGATTAGGCGTAGTCGGGCACGTCGTAGGGG (C0609)

Truncated sAC from *Salmon salar* (Met 1 to Ser 499) was constructed by successively PCRs using 5’ primer C3978 and 3’ primers C4256, C4257, C3682. Primer C3680 matches to the ApsAC to L577 and add a part of the HA-tag, primer C3681 adds part of the HA-tag, followed by a short linker (Gly, Ser, Gly), primer C3682 adds a hexa His-tag, followed by a stop codon and a XbaI site. The resulting PCR fragment was cloned into vector pcDNA6/V5-HisA (Invitrogen, Carlsbad, USA). The primer sequences were:

TAAGCTTCCACCATGGGCTGGATCAAGGGCGACGGCGAGATCGAG (C3978) GTCGGGCACGTCGTAGGGGTAGCTGTACACCTCGATCTCCTTCTCC (C4256) GTGGTGGCCGCTGCCGGCGTAGTCGGGCACGTCGTAGGGGTAGC (C4257) ATATCTAGATTAGTGGTGGTGGTGGTGGTGGCCGCTGCCGGCGTAG (C3682)

Truncated sAC from *R. norvegicus* with a HA-tag added to the C-terminal end representing the N-terminal domain (Met 1 to Val 469) of sAC (57) was cloned into pcDNA5/FRT (Invitrogen).

### Cloning, expression, and purification of the *Ap*sAC_t_ catalytic core

cDNA for *A. punctulata* sAC (aa 1-577) was subcloned into pQE70-2Z.4 (from U. Kutay, ETH Zürich), with the structure Z-Z-His6-TEV-Ap-sAC, via BamHI and HindIII sites. The solubility-enhancing Z-tag corresponds to the IgG-binding domain of *S. aureus* protein A (UniProt P02976, aa 158-269). *E. coli* M15 (Qiagen) was transformed with *Ap*sAC (1-577):pQE70-2Z.4 and grown in LB media to OD600 ≈0.7. After IPTG induction O/N at 20 °C, cells were harvested, chemically lysed and the cleared lysate purified via Ni-IMAC, TEV cleavage, reverse IMAC, and gel filtration on Superdex 200. The *Ap*sAC catalytic core corresponds to that of rat and human sAC and is very similar to the truncated alternatively splice form sAC_t_ of mammalian sAC. For simplicity, we refer to all catalytic-core constructs as sAC_t_.

The activity of purified *Ap*sAC_t_ protein was measured as follows. Duplicates of 100 µl containing 2 ng/µl purified *Ap*sAC_t_ (1-577) were incubated at 25 °C in a buffer containing (in mM): 100 BIS-TRIS-Propane at pH 7.0 to 8.0 in 0.1 pH steps, 12 NaCl, 120 KCl, 5 MnCl_2_, 1 MgCl_2_, 2 CaCl_2_ and 5 ATP. Buffer without protein at pH = 7.0 served as negative control. Aliquots (20 µl) were removed at 10-min intervals from 0 to 30 mins, quenched with 0.5% trifluoroacetic acid (TFA) and immediately shock-frozen in liquid N_2_. Upon assay completion, samples were thawed, spun for 5’ at 14,680 rpm, and 2 µl diluted with 100 µl MPW. The diluted samples were analyzed over a Prototype XBridge BEH C18 column (2.5 µm XP, 2.1 × 75 mm) at 30 °C with a gradient 2-100% ACN in 10 mM NH_4_OAc pH 4.5 over a 5’ run on a Waters Acquity UPLC. Peaks were integrated and cAMP signals normalised to % of total area (ATP + ADP +AMP + cAMP).

### Cell lines

For heterologous expression, HEK293 cells from the European Collection of Authenticated Cell Cultures (ECACC) were used. Mycoplasma tests were carried out regularly once a year. The results of the mycoplasma tests are available upon request.

### Generation of HEK293 cell lines stably expressing *Ap*sAC or *Ss*sAC or *Rn*sAC_t_

HEK293 cells were electroporated with plasmids using the Neon 100 Kit (Invitrogen, Carlsbad, USA) and a MicroPorator (Digital Bio) according to the manufacturer’s protocol (3 × 1245 mV pulses with a 10-ms pulse width). Cells were transferred into complete medium composed of M10 plus GlutaMAX or DH10 plus GlutaMax (Invitrogen) and 10% fetal bovine serum (Biochrom, Berlin, Germany). To select monoclonal cells stably expressing sAC, the respective antibiotic was added to the cell culture medium 24 h after electroporation. Monoclonal cell lines were identified by immunocytochemistry using a rat anti-HA antibody (Roche Applied Science). HEK293 cells stably overexpressing rat sAC_t_ were as described (58).

### Cyclase activity assay of recombinant human sAC_t_ protein

*In vitro* activity of GST-tagged *Hs*sAC_t_ was performed as described previously (15). Briefly, activity was assayed in 100 µl reactions containing (mM): 5 MgCl_2_, 5 CaCl_2_, 40 NaHCO_3_, 1 ATP, 50 Tris pH 7.5, and the indicated concentration of TDI-8164. Each reaction contained ∼1,000,000 counts of α-^32^P labeled ATP. The cAMP was purified using sequential Dowex and Alumina chromatography as previously described (59).

### Preparation of sperm samples and flagella

Collection of dry sperm from *Arbacia punctulata* was described previously (27). Briefly, 0.2– 0.5 ml of 0.5 M KCl was injected into the sea urchin cavity to induce spawning. Spawned sperm (dry sperm) were collected using a Pasteur pipette and stored on ice. Sperm from mature salmon males (*Salmon salar*) were provided by the Department of Fish Ecology of LANUV (State Agency for Nature, Environment, and Consumer Protection of the State of North-Rhine Westphalia) (Kirchhundem-Albaum, Germany) and the Aqua-Wild Salmon Centre (Siegburg, Germany). Briefly, mature males were anaesthetized with Tricain (MS222). Fish were rinsed and blotted clean using paper towels. Sperm samples were released by gentle abdominal pressure and collected in 100-ml PP cups with screw caps (Sarstedt, Nümbrecht, Germany) and immediately stored on ice. Stripped fish were returned to recovery tanks. Sperm density was calculated using a Neubauer counting chamber (Carl Roth; Karlsruhe, Germany). For counting, sperm cells were diluted 1:32,000 in PBS. Sperm density was 1.64⨯10^10^ ± 0.47⨯10^10^ cells/ml (n = 9). Preparation of sperm flagella were as described (9).

### Western blotting

Flagella and heads from *A. punctulata* sperm were prepared as described (9). Briefly, flagella were resuspended in solubilization buffer containing (mM): 140 NaCl, 1 EDTA, 10 Tris–HCl at pH 7.6, 1% dodecyl-maltopyranoside (DDM), and protease inhibitor cocktail 1:500 (P8340, Sigma Aldrich, USA)), incubated on ice for 60 min and sonicated three times for 30 s (Sonifier 450, Branson, Danbury, CT). After incubation, supernatant and pellet were separated by centrifugation (10,000×g at 4°C). Supernatant was subjected to Western blotting. Non-transfected HEK293 (control) and HEK293 cells stably expressing HA-tagged sAC were lysed by sonification in a hypotonic buffer containing (mM): 2 Hepes/NaOH at pH 7.4, 0.2 EDTA, 10 NaCl and 1:500 protease inhibitor mixture mPIC (Sigma Aldrich, St. Louis, USA). The suspension was subjected to Western blotting. Protein concentration was determined using the Bradford assay or the BCA test kit (Pierce) according to the manufacturer’s protocol. Solubilized sperm flagella and heads from *A. punctulata*, hypotonically treated HEK293 cells heterologously expressing HA-tagged sAC, and HEK293 control cells were separated by SDS-PAGE using TruPAGE precast gels (4-12%) (Sigma-Aldrich) or 8% Laemmli SDS-gels. Samples were denatured for 10 min at 95 °C prior to separation. Protein Marker VI (AppliChem, Darmstadt, Germany) was used as molecular weight markers. Proteins were transferred onto an Immobilon FL PVDF membrane (Merck Millipore, Darmstadt, Germany), probed with antibodies, and analyzed using the Odyssey Imaging System (LI-COR, Bad Homburg, Germany). Figures were prepared using CorelDraw and Photo-Paint software (both from Corel Corporation). Primary antibodies were: rat-anti-HA (1:5,000; Roche Applied Science catalog, Penzberg, Germany), mouse-anti-HA (1:5,000; Sigma-Aldrich). Secondary antibodies were IRDye680 and IRDye800 antibodies (LI-COR, 1:25,000).

### Measurement of sAC activity and quantification of cAMP content in sperm and cell lines

The cAMP synthesis of intact sperm from sea urchin and salmon under various salt and pH conditions on the sub-second time scale was analyzed using the stopped-flow instrument. Experiments on the seconds to minutes time scale were performed at room temperature in 96-well Protein LoBind plates (Eppendorf, Hamburg, Germany).

The cAMP synthesis of heterologously expressed sAC in HEK293 lysates was induced by applying di-sodium-ATP (Sigma Aldrich) and MgCl_2_ or MnCl_2_ in isotonic HEPES buffer. If not otherwise stated, reaction buffer contained (in mM): 1 ATP, 5 MgCl_2_ or 5 MnCl_2_, 0.3 CaCl_2_, 120 KCl, 12 NaCl, 0.2 IBMX, 50 HEPES/KOH (at pH 7 or pH 8), and optional 25 HCO_3_. Reaction time was 10 minutes for assays with HEK293 lysates.

For cAMP quantification using the catch-point assay, reactions were stopped by quenching with 0.5 × reaction volume HClO_4_ (1.5 M). The solution was neutralized with 0.625 × reaction volume K_3_PO_4_ (1 M). Salt precipitate and cell debris were removed by centrifugation of the 96-well plates (10 min, 4,000×g, 4 °C). The supernatant was transferred into a new 96-well plate.

The total content of cAMP in quenched samples was determined by using the CatchPoint cAMP Fluorescent Assay Kit (Molecular Devices, San Jose, USA) with some modifications, in particular to adjust reactions volumes to the 8- or 12-channel pipettes (Eppendorf, Hamburg, Germany), repetitive pipette HandyStep (Brand, Wertheim, Germany), or the Multi-Channel Auto Sampling System NSP-7000 (Nichiryo, Japan). Briefly, 60 µl of the quenched sample and 50 µl each of the anti-cAMP antibody and the HRP-cAMP conjugate were applied to an anti-rabbit IGG-coated 96-well plate according to the manufacturer’s protocol. After 90 min of incubation at room temperature and repetitive washing (3 times) with wash buffer (125 µl each), the Stoplight Red Substrate (125 µl) was applied to monitor the immunoreaction. The 96-well plates were analyzed using a FLUOstar Omega microplate reader (BMGLabtech, Ortenberg, Germany). The amount of cAMP was quantified using calibration curves obtained by serial dilutions of cAMP standards. Analysis was done using the reader’s data analysis software MARS. Further data processing was done using Excel (Microsoft, USA) and OriginPro (Origin lab, USA). Figures were prepared using CorelDrawX6 and Photo-PaintX6 software (Corel Corporation).

The cAMP content of intact HEK293 cells stably expressing *Rn*sAC_t_ (2.0 ⨯ 10^6^ cells) was determined by an accumulation assay (58). Cells were transferred to 1.5 ml tubes and incubated in DMEM + 10% FBS in suspension at 37°C, 5% CO_2_ for one hour. A time zero value for each condition was determined by adding 100 µl of cells directly into 100 μl stop solution (0.2 M HCl). To measure cAMP accumulation, cells in suspension were incubated for the indicated period of time in the presence of 500 μM IBMX and 30 µM TDI-8164 at 37°C after which 100 μl of cells were transferred to a fresh tube containing stop solution. Intracellular cAMP content was determined using Correlate-EIA Direct Assay (Assay Designs, Inc).

### Measurement of changes in V_m_, pH_i_, and [Ca^2+^]_i_, in *A. punctulata* sperm

We measured changes in Ca^2+^, pH_i_, and V_m_ by loading sperm samples with the corresponding dyes (Fluo-4AM, BCECF-AM, or pHrodo Red-AM from Molecular Probes, Eugene, USA, and FluoVolt from Thermofisher) and subsequent mixing in a rapid-mixing device (SFM-400, BioLogic, Claix, France) (27). Dry sperm was suspended 1:6 (v/v) in artificial seawater (ASW) containing (in mM): NaCl 423, CaCl_2_ 9.27, KCl 9, MgCl_2_ 22.94, MgSO_4_ 25.5, EDTA 0.1, HEPES 10 at pH 7.8, and the respective dye (18 °C). Loading concentrations and times were: Fluo-4AM: 10 μM for 30-45 min (0.5% Pluronic F127, Molecular Probes); BCECF-AM: 10 μM for 10 min; pHrodo Red-AM: 10 μM for 30-45 min (0.5% Pluronic F127). After 1:20 (v/v) dilution in ASW, sperm were allowed to equilibrate for 5 min. Subsequently, sperm were mixed 1:1 (v/v) with ASW. The 1:20 dilution experiments were performed by incubating sperm with the respective indicator in ASF (1:6). Sperm were then loaded into the stopped-flow device and rapidly mixed (1:20 dilution) with either ASF or ASW. The 1:2-dilution experiments were performed by incubating the sperm with the respective indicator in ASF (1:6), followed by 1:20 dilution in ASF. The diluted sperm suspension was then loaded into the stopped-flow device and rapidly mixed 1:2 with either ASF, 0KASW7.8 (ASW with no K^+^ present), or 30KASW7.8 (ASW with 30 mM K^+^). When mixing with 0KASW7.8, the final [K^+^] was 10 mM, i.e. similar to normal ASW; the final pH_o_ was about 7.75. When mixing with 30KASW7.8, the final [K^+^] was 30 mM and the final pH_o_ was 7.75. All signals obtained upon dilution of 1:2 and 1:20 in the stopped-flow were recorded at flow rates of 2 ml s^-1^.

Fluorescence was excited by pulsed LED light (SpectraX Light Engine, Lumencor, Beaverton, USA or M490L3, Thorlabs, Newton, USA) with a frequency of 10 kHz. Emission was recorded by photo-multiplier modules (H9656-20 and C7169, Hamamatsu Photonics, Japan). The signal was amplified and filtered by a lock-in amplifier (7230 DualPhase, Ametek, Paoli, USA). Data acquisition was performed with a data acquisition pad (PCI-6221, National Instruments, Austin, USA) and Biokine Software v.4.49 (BioLogic). Fluo-4 was excited using 490/20 nm filter (Semrock) and its emission collected using a 536/40 nm filter (Semrock); BCECF was excited using a 452/45 nm filter (Semrock). BCECF fluorescence was recorded in dual-emission mode using Brightline 494/20 nm and 540/15 nm filters (Semrock). The pH_i_ signals represent the ratio of F494/F540. All signals are the average of at least two recordings and are depicted as the percent change in ratio (ΔR/R) with respect to the first 10–20 data points after mixing. The pHrodo Red dye was excited at 572/15 nm and the emission was collected at 628/40 nm. Fluovolt was excited using a 513/18 nm filter (Semrock) and its emission collected at 540/20 nm (Semrock). Di-8-ANEPPS was exited at 475/20 nm (Semrock) and its fluorescence was recorded in dual-emission mode using Brightline 536/40 nm and 628/40 nm filters (Semrock). The V_m_ signals (ratio 536/549 nm (R); average of at least four recordings) are depicted as the percent change in ratio (ΔR/R) with respect to the first 10 data points. The signal recorded upon mixing sperm with ASF represented the baseline control and was subtracted from the respective signals. To manipulate cGMP in sperm during chemoattractant stimulation, sperm were incubated with 15 μM DEACM-caged cGMP for 7 min in the presence of 0.25% Pluronic™ F-127 (ThermoFisher). Uncaging was performed in the stopped-flow cuvette by a 360 nm LED (Thorlabs) with arbitrary waveforms (27).

### Measurements of pH_i_ in different compartments of single sperm cells

Sperm (1:25 dilution) were incubated for 30 min at 20°C in ASW containing 20 µM pHrodo Red-AM (Molecular Probes) and 0.5% Pluronic F127 (Sigma-Aldrich). After labelling, sperm were diluted (1:100 dilution) and loaded into glass capillaries for perfusion (Rectangle Boro Tubing 0.2x 4 mm; Vitro Tubes). pH signals were calibrated using the pH_i_ “pseudo-null point” method as previously described (27) using different ratios of a weak acid (propionic acid; pK_a_ = 4.88) and a weak base (ammonium chloride; pK_a_ = 9.24). pHrodo Red fluorescence was recorded with an inverted microscope (Olympus IX71) equipped with a 20x objective lens (UPlanSApo 20x, 0.75 NA; Olympus) and a 560 DCXR dichroic mirror (Chroma). Stroboscopic imaging of the rapid flagellum was achieved using a multicolor lamp (2 ms light pulses; Spectra X Light Engine; Lumencor). To accommodate for the large difference of emission intensity between head and flagellum, we used two alternating excitation wavelengths and excitation band-pass filers. Green excitation (ET545/30; Chroma) was used to image the sperm flagellum and Teal excitation (FF01-513/17; Chroma) for imaging of the head. Emission light was filtered using a band-pass filter (BA575-625; Olympus) and images were collected using an EMCCD camera (DU-897D; Andor Technology). Quantification of signals was done using custom-made software written in Matlab (Mathworks). Sperm cells were perfused with different null-point solutions, and the fluorescence intensity was recorded. pHrodo Red intensity was averaged for 30 frames before and after perfusion with the null-point solutions. To evaluate the resting pH_i_, a least-mean-square method was used. For this, individual points were weighted by the inverse of their s.e.m.

### Emulation of spawning in the stopped-flow device

Spawning was emulated in the stopped-flow device by two different approaches. Dry sperm was incubated 1:6 in artificial seminal fluid (ASF), containing the respective indicator dyes. ASF was ASW at pH 6.7 containing 30 mM [K^+^]. After incubation, the sperm suspension was diluted 1:20 in ASF. In the stopped-flow device the suspension was rapidly mixed with K^+^-free ASW that was fortified with 50 mM Hepes at pH 7.8. After 1:2 mixing the [K^+^] was 10 mM and pH was 7.7-7.8. Alternatively, dry sperm was incubated 1:6 in ASF and then mixed 1:40 with ASW in the stopped-flow device.

### The pH-clamp method

We determined the pH_i_ sensitivity of the alkaline-induced Ca^2+^ influx using the “pH_i_ pseudo-null-point” method (34, 60-62) (Bond and Varley, 2005; Chow et al., 1996; Eisner et al., 1989; Swietach et al., 2010) that allows clamping of pH_i_ to fixed values and calibration of the pH indicator BCECF. The method for sea urchin sperm is described in (24, 27). Key is a set of pH_i_-clamp solutions composed of a weak acid (butyric acid, BA) and a weak base (trimethylamine, TMA) at different molar ratios. TMA and BA freely equilibrate across the membrane and, at sufficiently high concentrations, establish a defined pH_i_ that is set by the acid/base ratio (61). The pH_i_-null-point solutions were prepared according to the following equation: pH_i_-null = pH_o_ – 0.5 log ([BA]/[TMA]); pH_o_ = extracellular pH (7.8) (34), wherein [TMA] indicates the concentration of trimethylamine and [BA] that of butyric acid. The [BA] was 15 mM.

### Measurement of sperm motility

Sperm motility was recorded in an inverted microscope (Olympus IX71) equipped with a 10x objective lens (UPlanSApo 10x, 0.4 NA; Olympus) under dark-field illumination at 25 Hz (sea urchin sperm) or 40 Hz (salmon sperm) with an EMCCD camera (DU-897D; Andor Technology). *Arbacia punctulata* sperm were diluted 1:200 in artificial seminal fluid (ASF) containing 0.5% Pluronic F127 (Sigma-Aldrich) and loaded into glass capillaries for superfusion (Rectangle Boro Tubing 0.2×4 mm; Vitro Tubes). The 150-mm loading tube (0.86 mm ID, 1.52 mm OD Fine Bore Polythene Tubing; Smiths medical) was then placed into artificial seawater (ASW) with 0.5% Pluronic F127 to prevent sticking to the glass surface. Starting the perfusion results in mixing ASF with sperm and replacing the ASF with ASW. After dilution of the sperm suspension by 1:100 in ASW, the perfusion was stopped and the time course of activation was recorded. Quantification of the number of activated cells at different time points was done using custom-made CASA software written in MATLAB (Mathworks).

*Salmon salar* sperm (dilution 1:2770) were loaded for five minutes in salmon ASF containing 4 µM DEACM-caged cAMP and 0.5% Pluronic F127 (Sigma-Aldrich). Incubations were discarded after 30 minutes. For imaging, sperm were loaded into custom-made glass chambers with 150 µm height. A 390-nm light source (SpectraX Light Engine; Lumencor) was used to release cAMP. The power of the UV (147 mW maximal power) was measured using a power meter (Controller PowerMax and head model PS19Q; Coherent). The flash intensity was graded using neutral density filters (Absorptive ND Filters; Thorlabs), and the duration of the flash (5 ms length) was controlled using a custom-made LabVIEW software. The percentage of cAMP released was calculated based on the photochemical properties of DEACM-caged cAMP (63) and the light intensity at the focal plane of the microscope. The motility indicator was calculated using custom-made software written in MATLAB (Mathworks). In brief, the software binarizes the images using an automatic threshold (graythres). Sperm motility can be coarsely quantified by subtracting temporally-adjacent frames. The absolute value of this image difference is then integrated to produce a motility score. When sperm are perfectly static, such score becomes zero and gradually increases as cells move. The relative sperm density was quantified as the mean pixel intensity of the binarized image. To compensate for slight variations in sperm density, this value was used to normalize the motility score.

## Data Availability

All relevant data are available from the authors upon request.

