## Supplementary Information for "External fertilization is orchestrated by a pH-regulated soluble adenylyl cyclase controlling sperm motility and chemotaxis"

### Supplementary Figures and Tables

#### Supplementary Table 1: Functional amino acid residues in sAC[a]

| position | function |
|---|---|
| D47 | ATP/ion binding site |
| **K95**[b] | anchoring site of $HCO_3^-$ binding |
| D99 | ATP/ion binding site |
| T405 | interacts with the 6-$NH_2$ group of ATP |
| **R176** | key switch between catalytic and regulatory ($HCO_3^-$) sites |
| K144 | interacts with $\beta P_i$ group |
| R416 | interacts with $\alpha P_i$ group |
| D339 | 2/3 beta loop around D339, interacts with R176 |
| F338 | flips over from near R176 into a M419/M300 cleft after bicarbonate binding |
| K334 | binds to N1 of the adenine ring of ATP |

[a]numbers refer to positions in the human sAC sequence (Kleinboelting et al. 2014)

[b]non-conserved residues are in bold.

**Supplementary Table 2: Key amino-acid residues in the $HCO_3^-$-binding site.**

| Phylum | Common name | Species | | |
|---|---|---|---|---|
| **Chordata** | **Mammals** | ***Homo sapiens*** | **K95** | **R176** |
| Dinoflagellata | Algae | *Symbiodinium microadriaticum* | K | S |
| Echinodermata | Sea urchin | *Strongylocentrotus purpuratus* | K | N |
| | | *Arbacia punctulata* | K | N |
| | Starfish | *Acanthaster planci* | K | N |
| Porifera | Sponge | *Amphimedon queenslandica* | K | T |
| Priapulida | Marine worms | *Priapulus caudatus* | N | R |
| Mollusca | Oysters | *Crassostrea gigas* | K | N |
| | Scallop | *Patinopecten yessoensis* | K | N |
| Brachiopoda | Lamp shell | *Lingula anatina* | K | N |
| Cnidaria | Corals | *Acropora danai* | K | N |
| | | *Pocillopora damicornis* | K | N |
| | Sea anemone | *Nematostella vectensis* | K | N |
| | | *Exaiptasia pallida* | K | N |
| Hemichordata | Acorn worm | *Saccoglossus kowalevskii* | K | R |
| | Spring tail | *Folsomia candida* | K | S |
| Arthropoda | Insect | *Aedes albopictus* | K | S |
| Chordata | Sea squirt | *Ciona intestinalis* | K | N |
| | Lancelet | *Branchiostoma belcheri* | - | N |
| | | *Branchiostoma floridae* | K | N |
| | Gekko | *Gekko japonicus* | N | R |
| | Teleost fish | *Danio rerio* | - | - |
| | | *Salmo salar* | N | R |
| | | *Clupea harengus* | N | R |
| | | *Esox lucius* | N | R |
| | | *Oncorhynchus mykiss* | N | R |
| | | *Pygocentrus nattereri* | N | R |
| | | *Latimeria chalumnae* | K | R |
| | Cartilaginous fish | *Squalus acanthias* | N | R |
| | | *Rhincodon typus* | N | R |
| | | *Callorhinchus milii* | N | R |
| | Tortoise/turtle | *Chelonia mydas* | N | R |
| | | *Pelodiscus sinensis* | T | R |
| | | *Terrapene carolina mexicana* | N | R |
| | Alligator | *Alligator mississipiensis* | N | R |
| | | | N | R |
| | Crocodile | *Crocodylus porosus* | N | R |
| Bacteria | Cyanobacteria | *Arthrospira platensis* | K | R |

**Supplementary Table 3: Composition of seminal fluid of *A. punctulata* and *S. salar***

| | ***A. punctulata*[1]** | | ***S. salar*[1]** | |
|---|---|---|---|---|
| component | seminal fluid[2] | artificial seawater[3] | seminal fluid[4] | fresh water[5] |
| pH | 6.8 ± 0.25 | 7.8 | 8.3 | 8.1 |
| $Na^+$ | 430.8 ± 12.8 | 423 | 110 | 0.5 |
| $K^+$ | 25.7 ± 2.1 | 9 | 37 | 0.1 |
| $Ca^{2+}$ | 9.5 ± 0.2 | 9.3 | 0.65 | 0.8 |
| $Mg^{2+}$ | 51.1 ± 1.2 | 48.5 | 1.15 | 0.2 |
| $Cl^-$ | 514.9 ± 10.8 | 496.5 | 109 | 0.7 |
| osmolality | 1058.7 | 1011.8 | 265 | 3-4 |

[1]concentrations in mM, except pH; [2]ions: n = 6 experiments, pH: n = 5 experiments; [3](Hamzeh et al., 2019); [4](Rosengrave et al., 2009, *Comp Biochem Physiol* 152, 123-129); [5]Wahnbachtalsperrenverband Siegburg, Germany.

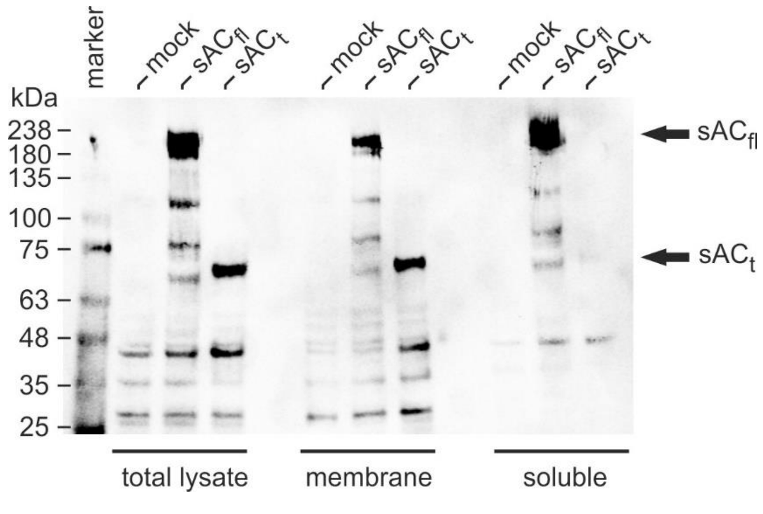


**Supplementary Figure 1**. Membrane association of *A. punctulata* $sAC_{fl}$ and $sAC_t$ heterologously expressed in HEK293 cells.

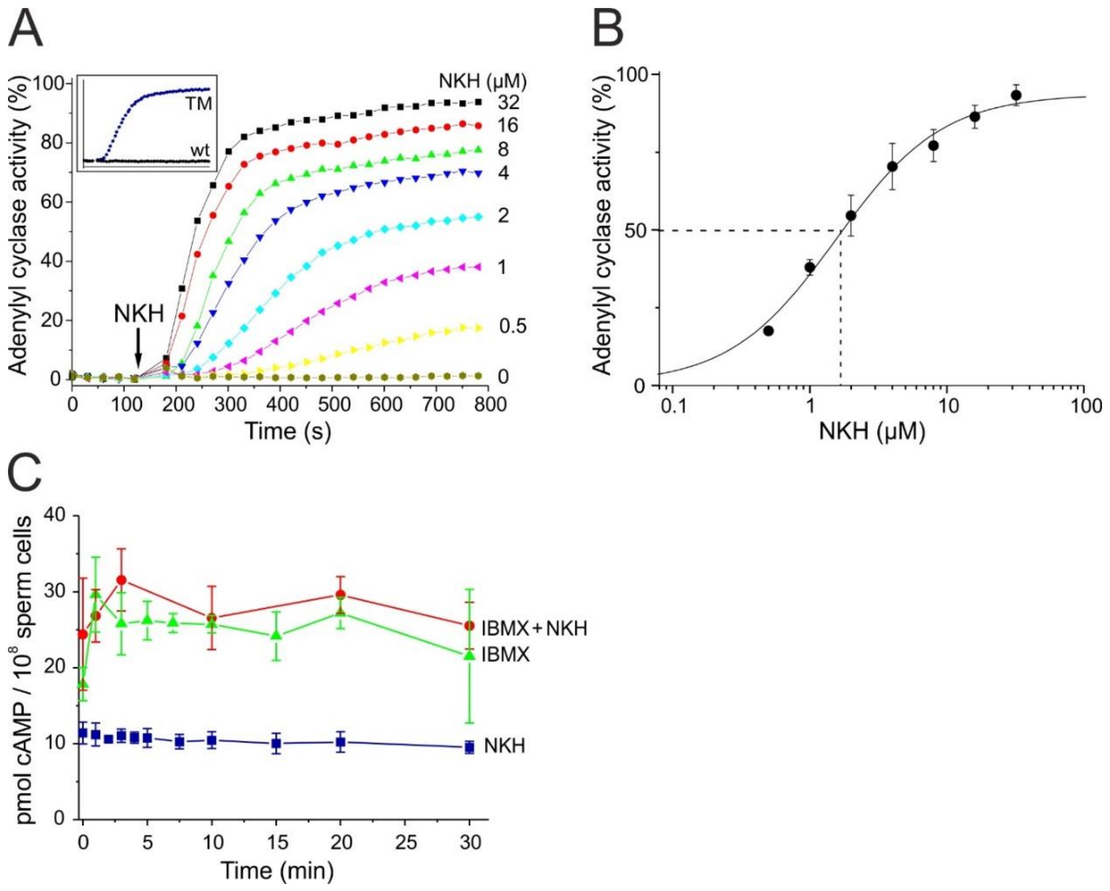


**Supplementary Figure 2**. **Dose-dependent changes in cAMP concentration in HEK 293 cells stimulated by NKH477.**

The changes in cellular cAMP concentration were monitored using Fluo-4-loaded HEK293 cells heterologously expressing a cyclic nucleotide-gated channel (CNGA2) as sensor for cAMP (TM). Non-transfected Fluo-4-loaded HEK293 cells (wt) were used as control (inset). **(A)** Kinetics of NKH477-induced changes in cAMP. NKH477 was added at t = 120 s. Inset: Fluorescence of CNGA2 cells and control cells at saturating NKH477 (30 μM) to demonstrate specificity of sensor activity **(B)** Dose-response relationship for NKH477 at t = 750 s. Mean ± s.d. (n = 3). **(C)** Changes in cAMP concentration after stimulation with NKH (40 μM, blue), NKH (40 μM) + IMBX (1 mM, red), and IBMX (1 mM, green); n = 3 animals, NKH doublets, NKH + IBMX doublets, IBMX for animal #1 doublets, for animals #2 and 3 single data. The cAMP concentration of unstimulated sperm is given at t = 0.

**Supplementary Information to Fig . S2**

**Transmembrane ACs do not contribute to cAMP synthesis in *A. punctulata* sperm.**

The tmAC isoforms AC1, 2, 5, and 9 reportedly exist in sea urchin sperm (7, 13) (Beltrán et al., 2007; Vacquier et al., 2014). However, their physiological significance is unclear because sAC is highly abundant (9) (Trötschel et al., 2020) and accounts for >95% of cAMP synthesis (7, 26) (Bookbinder et al., 1990; Vacquier et al., 2014). Nonetheless, we examined the contribution of tmACs to cAMP synthesis in intact *A. punctulata* sperm using NKH477, a water-soluble analogue of forskolin that potently activates tmACs (except rodent tmAC9) but not sAC (64) (Kamenetsky et al., 2006). As a positive control, we measured the action of NKH477 in HEK293 cells expressing a $Ca^{2+}$-permeable CNG channel as cAMP sensor (HEK-CNG) (65) (Wachten et al., 2006). Activation of tmACs by NKH477 faithfully elevated cAMP levels in HEK-CNG cells, as indicated by a $Ca^{2+}$ influx (Supplementary Fig. 2). By contrast, cAMP levels in sea urchin sperm were similar before and after stimulation with NKH477 (11.4 ± 1.5 pmoles cAMP/$10^8$ cells versus 9.5 ± 0.8 pmoles cAMP/$10^8$ cells (n = 3)) (Fig. 4A). It is noteworthy that IBMX, a broadly-specific inhibitor of phosphodiesterases (PDE), elevated cAMP levels in sperm, but application of IBMX and NKH477 simultaneously had no additional effect (Fig. 4A). Thus, *A. punctulata* sperm do not appear to contain significant levels of functional tmACs, suggesting that sAC is the major, if not only, source of cAMP synthesis.

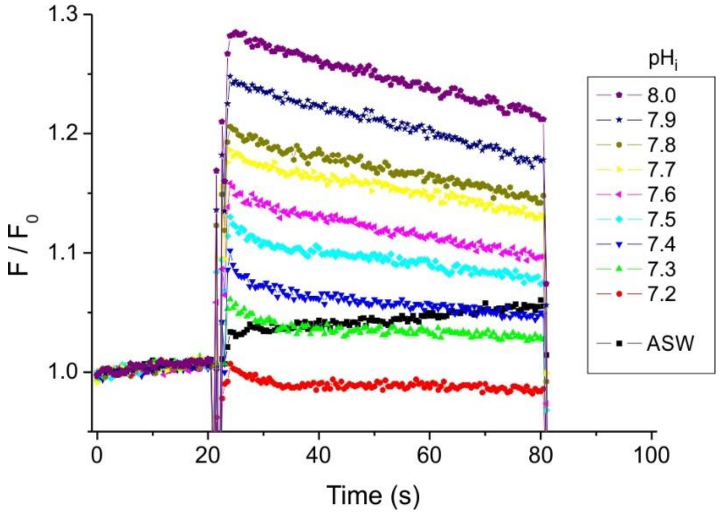


**Supplementary Figure 3.** Changes in intracellular $pH_i$ of *A. punctulata* sperm ($3 \cdot 10^8$ cells/ml) after addition of different pH-clamp solutions at t = 20 s as indicated by color code.

Changes in $pH_i$ were measured in a Fluostar device with the pH indicator dye BCECF (10 µM); $\lambda_{exc}$ = 485 nm, $\lambda_{em}$ = 520 nm.

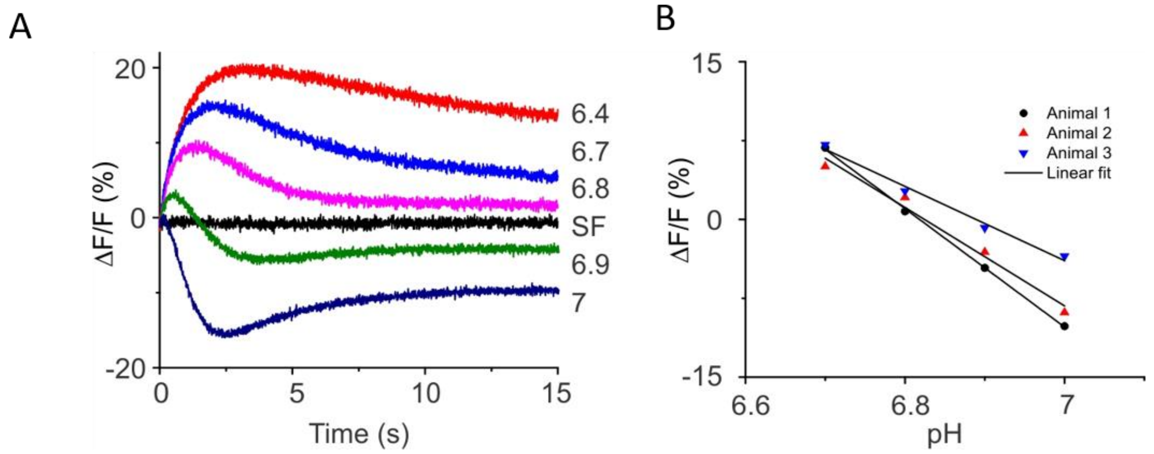


**Supplementary Figure 4. Determination of $pH_i$ of *A. punctulata* sperm by the pH-clamp null-point technique.**

(**A**) Changes in relative fluorescence ΔF/F (%) after mixing sperm with pH-clamp solutions in artificial seminal fluid (ASF). (**B**) Plot of ΔF/F (%) *versus* pH-clamp value for three animals. The mean pH value at ΔF/F = 0 was 6.84 ± 0.02 (n = 3).

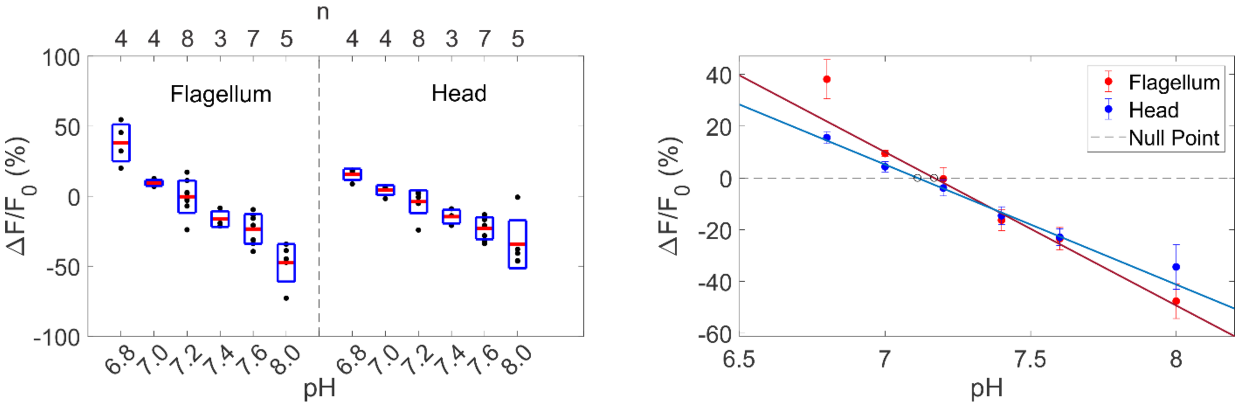


**Supplementary Figure 5. Resting $pH_i$ in different compartments of *A. punctulata* sperm.** pH signals from the sperm flagellum and head using the pH indicator pHrodo Red. Sperm cells were perfused with pH-clamp solutions that adjust $pH_i$ values from pH 6.8 to 8.0. Changes in relative fluorescence $\Delta F/F_o$ were quantified using fluorescence microscopy. *Left*: box plots of the changes in fluorescence upon perfusion. Red lines represent the average. Blue boxes represent the s.d.; values for individual sperm cells are shown as black dots. *Right*: linear fit of the changes in fluorescence versus $pH_i$ using the least-mean-square method. Interpolation to the null point ($\Delta F/F_o = 0$) yields a resting pH of 7.11 ± 0.01 for the head and 7.17 ± 0.01 for the flagellum. Error bars represent the s.e.m. Number of cells n are given on top of the left panel.

**Supplementary Movie 1.** Movie shows salmon sperm bathed in ASF under dark field illumination. At frame 100, a 5-ms flash of UV light was applied that produced 97 nM cAMP. The flash is indicated by white frames in the movie. Movie is displayed at original speed of 40 fps. Sperm were incubated with DEACM-caged cAMP (4 μM) for 30 min.
